# Conformational heterogeneity of the BTK PHTH domain drives multiple regulatory states

**DOI:** 10.1101/2023.06.02.543453

**Authors:** David Yin-wei Lin, Lauren E. Kueffer, Puneet Juneja, Thomas Wales, John R. Engen, Amy H. Andreotti

## Abstract

Full-length BTK has been refractory to structural analysis. The nearest full-length structure of BTK to date consists of the autoinhibited SH3-SH2-kinase core. Precisely how the BTK N-terminal domains (the Pleckstrin homology/Tec homology (PHTH) domain and proline-rich regions (PRR) contain linker) contribute to BTK regulation remains unclear. We have produced crystals of full-length BTK for the first time but despite efforts to stabilize the autoinhibited state, the diffraction data still reveals only the SH3-SH2-kinase core with no electron density visible for the PHTH-PRR segment. CryoEM data of full-length BTK, on the other hand, provide the first view of the PHTH domain within full-length BTK. CryoEM reconstructions support conformational heterogeneity in the PHTH-PRR region wherein the globular PHTH domain adopts a range of states arrayed around the autoinhibited SH3-SH2-kinase core. On the way to activation, disassembly of the SH3-SH2-kinase core opens a new autoinhibitory site on the kinase domain for PHTH domain binding that is ultimately released upon interaction of PHTH with PIP _3_. Membrane-induced dimerization activates BTK and we present here a crystal structure of an activation loop swapped BTK kinase domain dimer that likely represents the conformational state leading to trans-autophosphorylation. Together, these data provide the first structural elucidation of full-length BTK and allow a deeper understanding of allosteric control over the BTK kinase domain during distinct stages of activation.

## Introduction

The TEC family kinase, BTK (Bruton’s tyrosine kinase) is best known as the target of ibrutinib (IMBRUVICA®), the first-in-class covalent kinase active site inhibitor used to treat chronic lymphocytic leukemia (CLL), mantle cell lymphoma (MCL), Waldenström’s macroglobulinemia (WM), and chronic graft-versus-host disease (cGVHD). The TEC kinases are the second largest sub-family of non-receptor tyrosine kinases in the human genome after the SRC family [1-3]. The TEC and SRC kinases share the SH3-SH2-kinase domain arrangement (the ‘Src module’) (Fig. 1a). Structures of the autoinhibited SRC family kinases were solved in 1997 revealing, for the first time, the compact arrangement of the SH3 and SH2 domains assembled onto the distal side of the catalytic kinase domain [4]. Despite the shared ‘Src module’ of the SRC and TEC families, eighteen years passed before an autoinhibited structure of one of the TEC kinases was resolved by x-ray crystallography. In 2015 the BTK SH3-SH2-Kinase fragment was crystallized [5]. This BTK fragment crystallizes as a domain swapped dimer; the SH2 domain opens to allow the B helix of one chain to pack against the β-sheet of the other chain. One-half of the domain swapped BTK structure represents the current model for autoinhibited BTK and closely resembles the compact structure of autoinhibited SRC kinases (Fig. 1b).

**Figure 1.**
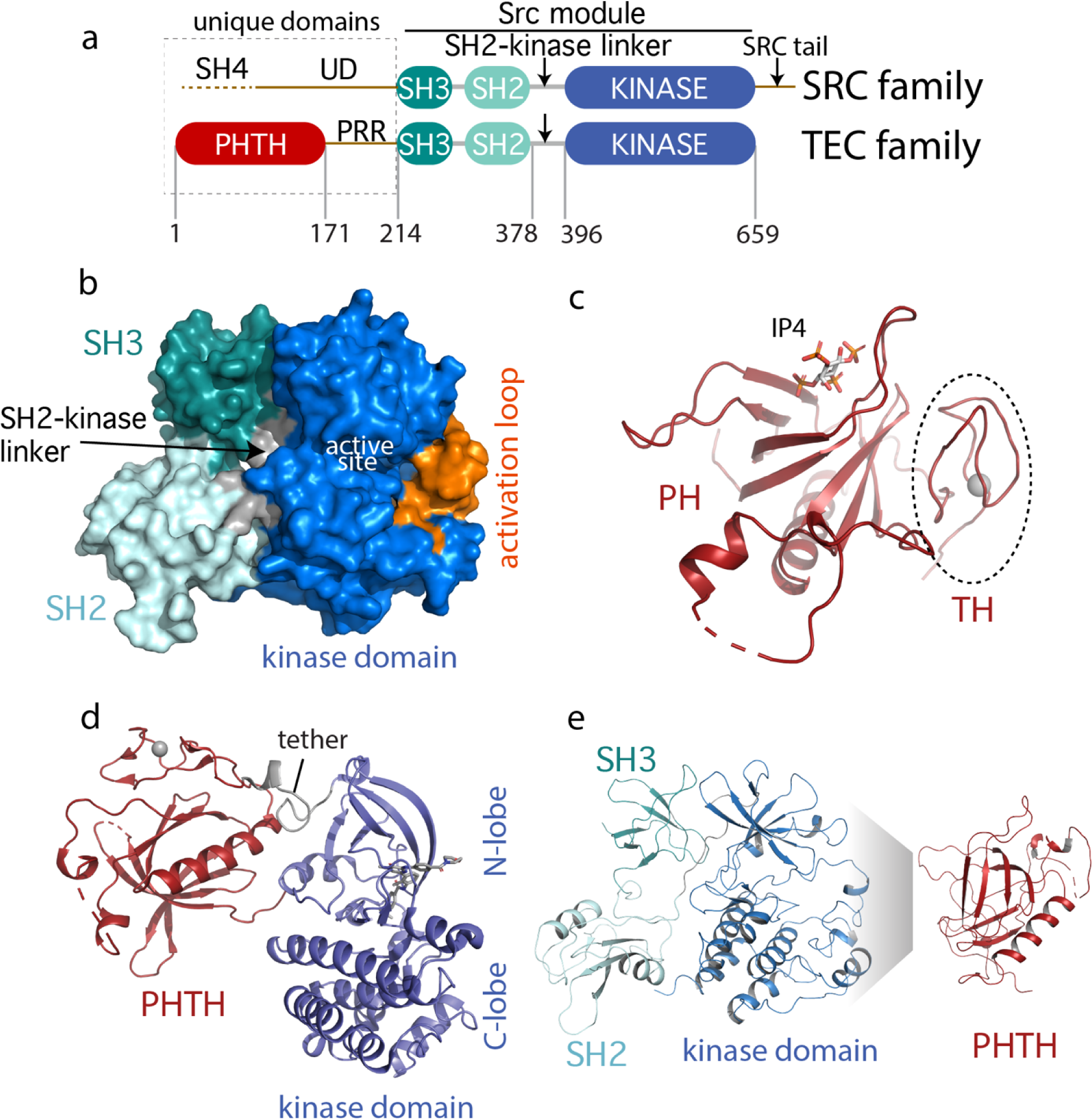
Current BTK structural data. (a) Comparison of the SRC and TEC domain architectures. Linker regions and domains are labeled, residue numbering shows BTK domain boundaries. The ‘Src module’ is the SH3-SH2-kinase region shred by both families. (b) Autoinhibited BTK core (Src module). The compact structure of the SH3-SH2-kinase region of BTK is exacted from the domain swapped dimer structure (PDB: 4XI2) solved by Wang et al. [5]. The three domains (SH3, SH2, kinase), the SH2-kinase linker, the activation loop and the active site are labeled. (c) Structure of the BTK PHTH domain bound to IP_4_ (PDB: 1B55) [8]. The monomer is shown for clarity and the TH region bound to Zn^2+^ is circled. (d) Structure of the tethered PHTH-kinase construct (PDB: 4Y93). (e) Solution based mapping of BTK PHTH interaction across the activation loop face of the kinase domain.

Outside of the Src module, the sequence and domain structures of the TEC and SRC kinases diverge. The SRC kinases contain a C-terminal tail (absent in the TEC family) that, upon phosphorylation on a conserved tyrosine, binds to the SH2 domain in an intramolecular fashion to stabilize the autoinhibited Src module. BTK instead stabilizes the SH2 autoinhibitory pose via a conserved acidic side chain at the end of the kinase domain [6]. The SRC kinases also contain an N-terminal SH4-unique domain (UD) region (Fig. 1a) that is intrinsically disordered and has not been resolved in the available crystal structures. The SRC kinases are myristoylated or palmitoylated in the SH4 domain driving membrane association [7]. In contrast, BTK contains an extended N-terminal region (PHTH-PRR) consisting of the phospholipid binding Pleckstrin Homology (PH) domain followed by a Tec Homology (TH) domain and a long linker housing two proline-rich regions (PRR) (Fig. 1a). The folded N-terminal PH domain of BTK has been crystallized [8] revealing a binding site for inositol 1,3,4,5-tetrakisphosphate (IP _4_), the headgroup of its target phosphatidylinositol (3,4,5)-trisphosphate (PIP _3_) ligand (Fig. 1c). The crystal structure shows the TH domain bound to zinc and closely associated with the PH domain; the folded N-terminal domain of BTK is hereafter referred to as PHTH. The BTK PHTH domain crystallizes as a homodimer, named for the preeminent structural biologist, Matti Saraste, who solved the first BTK PHTH structure [9].

The N-terminal PHTH-PRR domains account for over 30% of the BTK sequence. The BTK PHTH domain structure has been solved both as the isolated domain and in a construct where PHTH is covalently attached to the BTK kinase domain through a short tether [5]. The tightly tethered structure revealed an interface between the PHTH domain and the N-lobe of the kinase domain (Fig. 1d) but the contacts are mutually exclusive with the contacts between the BTK SH3 and kinase domain in the autoinhibited ‘Src module’ (Fig. 1b) raising questions about whether both the SH3 and PHTH domain associate with the kinase domain simultaneously. At the same time, we conducted solution-based NMR and hydrogen/deuterium exchange mass spectrometry (HDXMS) studies [10, 11] that point to autoinhibitory interactions between the PHTH domain and the activation loop face of the BTK kinase domain (Fig. 1e). In this case, the mapped interface reflects intermolecular contacts that result upon addition of excess PHTH domain to the isolated BTK kinase domain (NMR) or comparison of full-length BTK with the SH3-SH2-kinase fragment (HDXMS). Thus, the BTK structural data reported to date support two poses for the N-terminal PHTH domain: PHTH domain landing on the distal face of the kinase domain N-lobe (Fig. 1d) and PHTH domain binding to the activation loop face of the kinase domain (Fig. 1e).

Here, we set out to solve the structure of the full-length BTK protein. Our crystallization target included the N-terminal PHTH domain and the long PRR linker region followed by the SH3, SH2, SH2-kinase linker, and kinase domains (Fig. 1a). Mutations were introduced into the SH3-SH2-kinase core in an effort to stabilize the compact autoinhibited conformation. We reasoned that this approach might encourage the PHTH domain to adopt a stable autoinhibited state or states. Despite efforts to optimize the autoinhibitory conformation of BTK, the crystallography presented here reveals the same domain swapped dimer solved previously and no electron density is observed for the PHTH-linker residues. Further analyses using both small angle X-ray scattering (SAXS) and cryo-electron microscopy (cryo-EM) support a BTK autoinhibited state wherein the core SH3-SH2-kinase region adopts the compact monomeric form much like that observed in the early SRC kinase structures. CryoEM allows the PHTH domain to be directly observed; the globular PHTH adopts a range of conformational states that surround but do not contact the core ‘Src module’. To revisit the PHTH domain crystallographically, we lengthened and redesigned the tether between PHTH and kinase domains to provide greater conformational freedom between domains. The resulting crystal structure revealed yet another binding pose for the BTK PHTH domain on the BTK kinase domain and mutational analysis supports a regulatory role for this interface in the full-length kinase. Finally, we have also captured a structure of a BTK kinase domain dimer revealing a potential arrangement of this domain undergoing trans-autophosphorylation following PIP_3_ mediated dimerization. Together, the structural and biochemical data presented here provide new insights into the allosteric control of BTK catalytic function. These findings create new opportunities to target this kinase in the context of disease and/or drug resistance.

## Results

### Construct design and optimization

To stabilize the autoinhibited conformation of full-length BTK for crystallography, we first optimized the inhibitory contacts between the SH3 domain and the N-lobe of the BTK kinase domain. Following previous efforts that successfully stabilized this region in a related SRC family kinase [12], we substituted key residues within the SH2-linker sequence with proline (construct ‘4P’ in Fig. 2a). The catalytic activity of the full-length BTK 4P variant, measured by autophosphorylation on Y511 in the BTK activation loop, is reduced compared to wild-type BTK (Fig. 2b). In addition, the melting temperature (T _m_) of BTK 4P is 4°C higher than wild-type BTK consistent with stabilization of the overall fold (Fig. 2c, Fig. 2-Figure supplement 1). Next, we tested the effect of mutation of L390 to phenylalanine on the activity and stability of full-length BTK. L390 resides in the SH2-linker and has been shown to stabilize the hydrophobic stack (W421 and Y461) in the kinase domain N-lobe [13]. The aromatic phenylalanine is found in other members of the TEC family suggesting it is well tolerated in this position. We find the L390F mutation does not change the activity of full-length BTK but it does increase the T _m_ by 1.5°C compared to wild-type BTK (Fig. 2b,c and Fig. 2-Figure supplement 1). We therefore combined the 4P and L390F mutations (BTK 4P1F) and observe a further decrease in autophosphorylation activity and 4.5°C increase in T_m_ (Fig. 2b,c and Fig. 2-Figure supplement 1). Lastly, we compared the BTK 4P1F mutant to full-length BTK containing ITK activation loop residues (L542M S543T, V555T, R562K, S564A, P565S) instead of the wild type BTK sequence. We have previously show that incorporation of these ITK activation loop residues results in dampened loop dynamics that favor the autoinhibited conformation [14]. As well, previous BTK crystallography efforts showed that these changes in the activation loop are required for crystallization [5]. As expected, the activity of full-length BTK with the ITK loop substitution is low and we find that this sequence change also stabilizes the overall protein as the T _m_ is 2.5°C higher than wild-type BTK (Fig. 2b,c and Fig. 2-Figure supplement 1). These data suggest that targeted modifications within the SH2-linker region and activation loop of BTK result in stabilization of the autoinhibited conformation.

**Figure 2.**
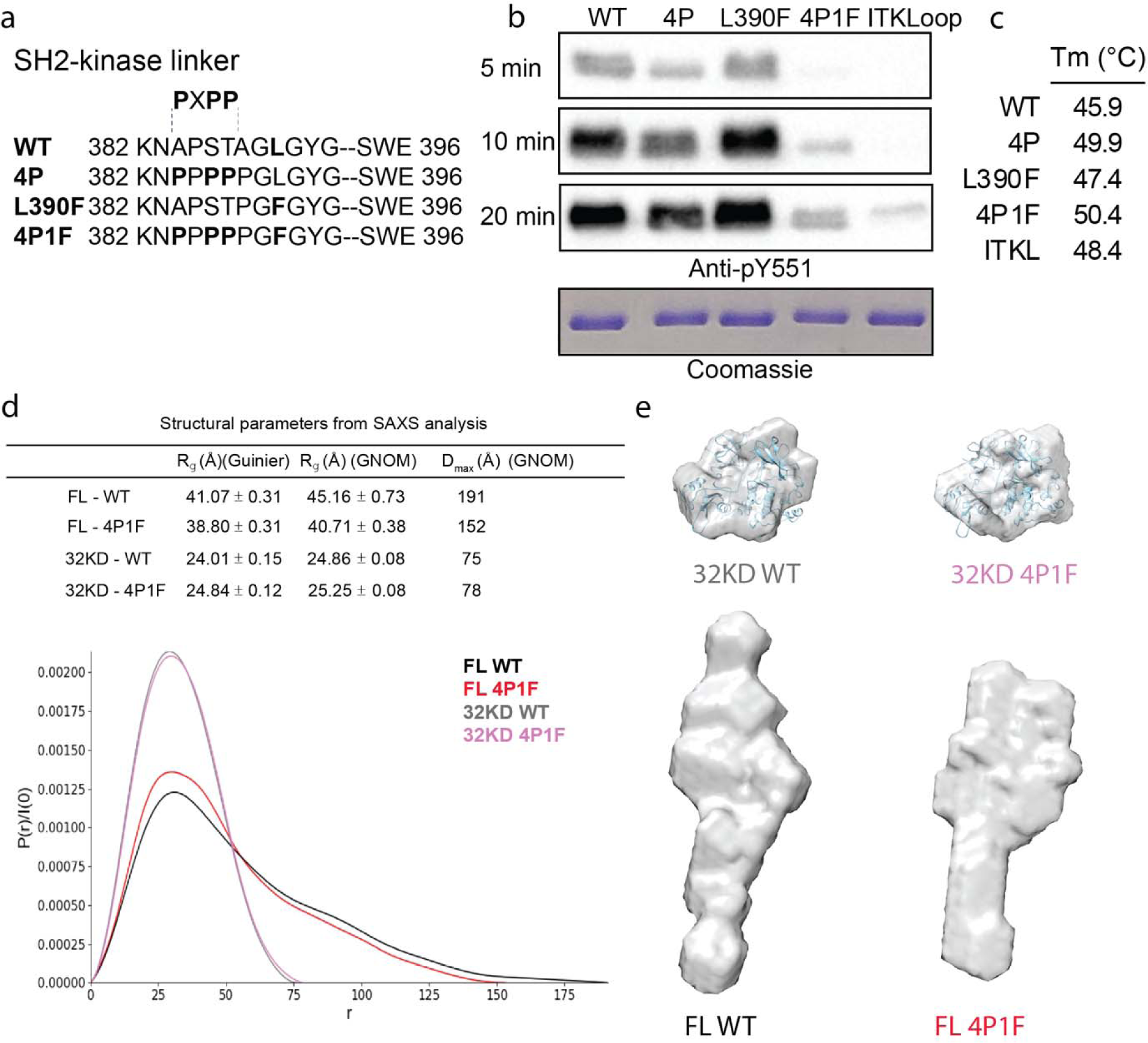
Stabilization of the BTK SH3-SH2-kinase core. (a) Mutations introduced into the SH2-kinase linker region of BTK (residues 382-396). PXPP indicates the region that adopts the left-handed, type II polyproline helix in the autoinhibited structure of BTK SH3-SH2-kinase. (b) Western blot showing the kinase activity of wildtype (WT) BTK, 4P, L390F, 4P1F, and ITKLoop BTK variants. Autophosphorylation on BTK is monitored using an anti-pY551 antibody and total protein levels are monitored by Coomassie stain. (c) Melting temperatures of BTK WT and variants (see Fig. 2-Figure supplement 1). (d) Distance distribution functions and structural parameters (R_g_ and D_max_) comparing the SH3-SH2-kinase fragment and full-length BTK (wildtype and 4P1F). (e) Surface representation of ab initio envelope reconstructions obtained from SAXS superimposed on the crystal structures for the BTK SH3-SH2-kinase fragment (top). Elongated envelopes for both full-length wild type (FL WT) BTK and the full-length 4P1F mutant of BTK are shown without structure superposition (bottom). Fig. 2-Figure supplement 1 provides Guinier and Kratky plots for all four BTK proteins. SASBDB accession codes are as follows: SASDRB9, SASDRC9, SASDRD9, SASDRE9.

Previous work on BTK made use of SAXS analysis to characterize both the full-length protein and domain fragments [15, 16]. Here we used SAXS to further characterize the effect of the sequence changes introduced to stabilize the autoinhibited state. Consistent with the previous observations [15, 16], the SH3-SH2-kinase fragment adopts a compact conformation while the full-length BTK protein is more extended (Fig. 2d,e). The SAXS derived structural parameters (R_g_ and D_max_) suggest that, compared to wildtype, the 4P1F BTK mutations have no effect on the average conformation of the SH3-SH2-kinase fragment. Ab initio shape reconstructions superimpose with the autoinhibited SH3-Sh2-kinase model derived from the previously solved structure (PDB: 4XI2) (Fig. 2e, top). In contrast, the volumes of both the ab initio envelopes of full-length wildtype and 4P1F BTK mutant are elongated (Fig. 2e, bottom). The full-length 4P1F mutant exhibits a somewhat more compact envelope (decreased particle dimension compared to wild-type full-length BTK) possibly consistent with the stabilized SH3-SH2-kinase core shifting the conformational preferences of the N-terminal PHTH-PRR region (Fig. 2d,e). Dimensionless Kratky and Grunier plots of the recombinant BTK proteins are provided in Fig. 2-Figure supplement 1.

With the stabilized full-length BTK mutant showing low activity (Fig. 2b) and a more compact conformational ensemble in solution (Fig. 2d,e), we built a full-length BTK construct for crystallization screens that incorporated the 4P1F mutations and the ITK loop (Fig. 3a). In addition, the catalytic K430 was mutated to arginine to eliminate kinase activity that could produce heterogeneity. This construct produced very small crystals in multiple crystallization conditions. To improve the size and quality of the crystals we introduced mutations at three side chains in the SH2 domain predicted to have high surface entropy [17]. The resulting construct, referred to as full-length BTK with stabilized core (FL BTKsc) (Fig. 3a), provided diffraction quality crystals (5-7 days at 4°C) both in the apo form and in the presence of various active site inhibitors. The highest resolution data collected (3.4 Å) was in the presence of the GDC-0853 (fenebrutinib) inhibitor.

**Figure 3.**
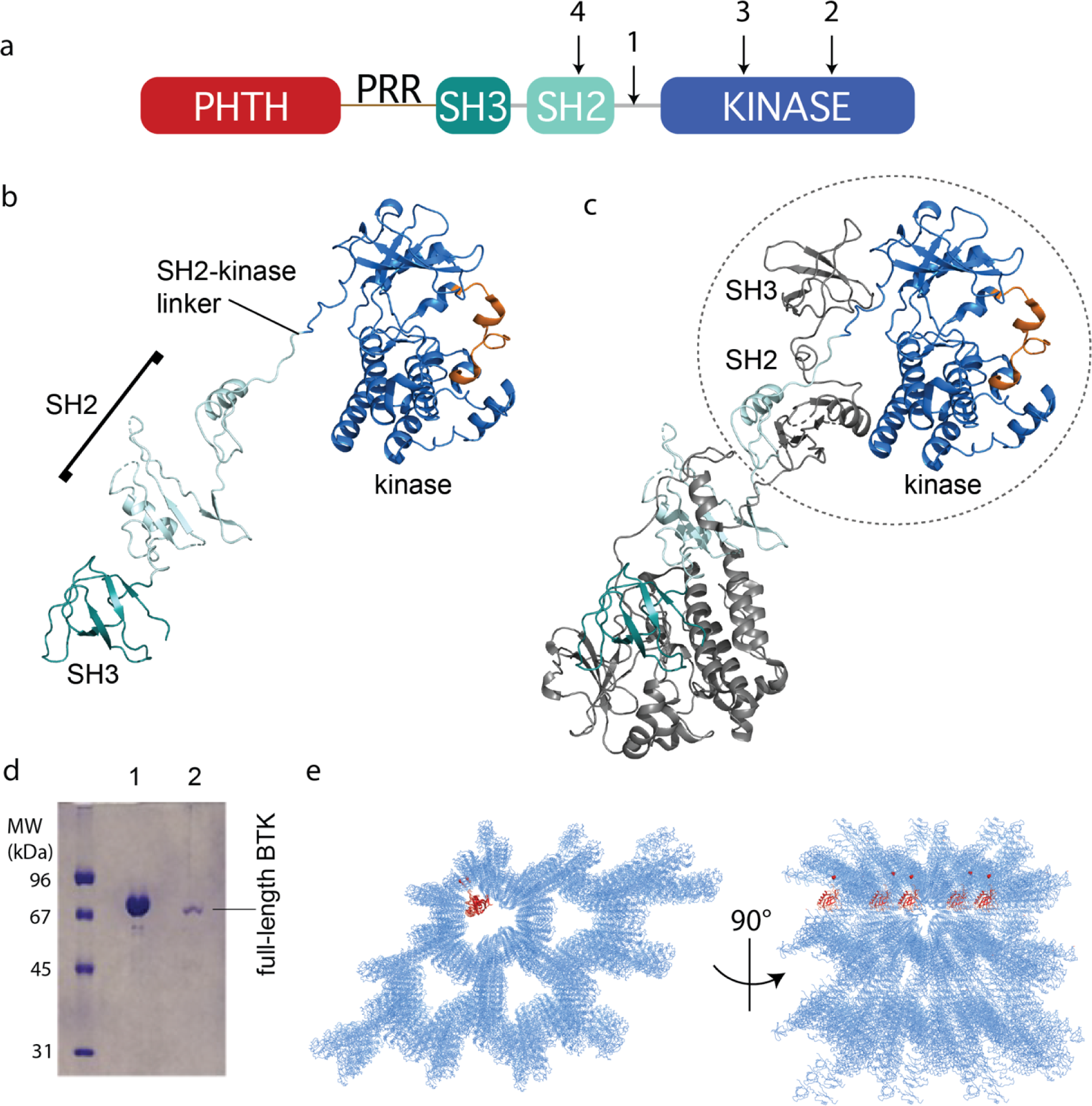
Crystallization of full-length BTK. (a) The crystallization target, full-length BTK with stabilized core (FL BTKsc), included (1) SH2-kinase linker mutations (4P1F: A384P, S386P, T387P, A388P, and L390F); (2) activation loop mutations (ITKLoop: L542M, S543T, V555T, R562K, S564A, P565S); (3) catalytic residue mutation (K430R); and (4) surface entropy reduction mutations (E298A, K300A, and E301A). The N-terminal domains, PHTH-PRR-SH3, are wildtype BTK sequence. (b,c) Structure of the BTK domain swapped dimer that results from crystallization of full-length BTK (PDB: 8GMB). PHTH-PRR region is missing from the electron density. One SH3-SH2-kinase monomer is shown in (b) and the autoinhibited SH3-SH2-kinase arrangement is circled in (c). Domain colors match those in (a) and the activation loop in the kinase domain is orange. (d) SDS-PAGE showing full-length BTK protein from crystals. Lane 1 is a purified full-length BTK control and lane 2 is protein derived from harvested and washed crystals. (e) Two views of crystal packing with the PHTH domain (red) modeled into one of the solvent channels.

## BTK crystal structure

Despite efforts to stabilize the autoinhibited form, the electron density derived from crystals of full-length BTK (FL BTKsc) defines the same domain-swapped dimer observed for the previously solved structure of the BTK SH3-SH2-kinase fragment [5] (Fig. 3b,c). A 16.9° rotation along residues 346-384 in the SH2 and SH2-kinase linker is observed for the domain swapped dimer structure solved here compared to that solved previously [5]. Notably, there is no PHTH domain density visible even after the structure is fully refined. To confirm that the crystals contain the full-length BTK protein and not a degradation fragment, we harvested, washed and analyzed the crystals by SDS-PAGE (Fig 3d). The result shows the crystal is full-length BTK protein with no evidence of smaller BTK fragments. Finally, we built five additional full-length BTK constructs with varying mutations/deletions within the long PRR containing linker between the PHTH and SH3 domains (see Materials and Methods). We find that four of the five constructs crystalize in a manner identical to FL BTKsc, which contains the wildtype sequence in the PHTH-PRR region, and no electron density is observed for the PHTH domain in any of these crystals (one construct did not form crystals).

The solvent content in the crystals of full-length BTK is high (67%) and crystal packing shows large solvent channels (Fig. 3e). Given the absence of PHTH domain electron density, we asked whether the previously solved PHTH domain structure could fit into the solvent channels. Placing the PHTH structure at the N-terminus of the BTK SH3-SH2-kinase model shows that the PHTH domain is readily accommodated within the solvent channels of the crystal (Fig. 3e). The crystallography results are consistent with a flexible N-terminal PHTH domain with the caveat that crystal packing might limit native autoinhibitory contacts between the PHTH and SH3-SH2-kinase regions. To address this, we examined the accessibility of the previously identified PHTH domain binding site on the activation loop face of the BTK kinase domain [11] in the context of symmetry related molecules in the crystal. The C-lobe G-helix portion of the PHTH/kinase interface mapped previously by solution methods [11], appears inaccessible while the activation loop itself and the portion of the N-lobe previously identified within the PHTH/kinase interface are accessible. Thus, it is possible that the PHTH domain mobility in the crystal is due to steric occlusion of the previously identified regulatory site. To further explore the domain arrangements within full-length BTK we next turned to single particle cryo-EM.

## CryoEM studies of full-length BTK

Using the same purified BTK protein used for structure determination by crystallography (FL BTKsc), we obtained cryoEM reconstructions of full-length BTK (Fig. 4). BTK particles were nicely distributed in the raw images (Fig. 4-Figure supplement 1). After several rounds of 2D classification, the 2D classes resembling the autoinhibited SH3-SH2-kinase core start to become visible (Fig. 4a,b, Fig. 4-Figure supplement 1). An extra unstructured density mass positioned adjacent to the main SH3-SH2-kinase core in addition to a globular density distant from the core are evident in the 2D classes and were persistently present after discarding specific 2D classes and repeated 2D classification. From ab-initio reconstruction and refinement, four 3D classes were obtained (Fig. 4-Figure supplement 1); three of which (Class 0, 1 and 3) were of sufficient quality to fit models of BTK SH3-SH2-kinase (Fig. 4c-e). The presence of the monomeric SH3-SH2-kinase region in all of the 3D reconstructions is significant as all BTK crystal structures to date have revealed a domain swapped dimer with two polypeptide chains contributing to the autoinhibited conformation.

**Figure 4.**
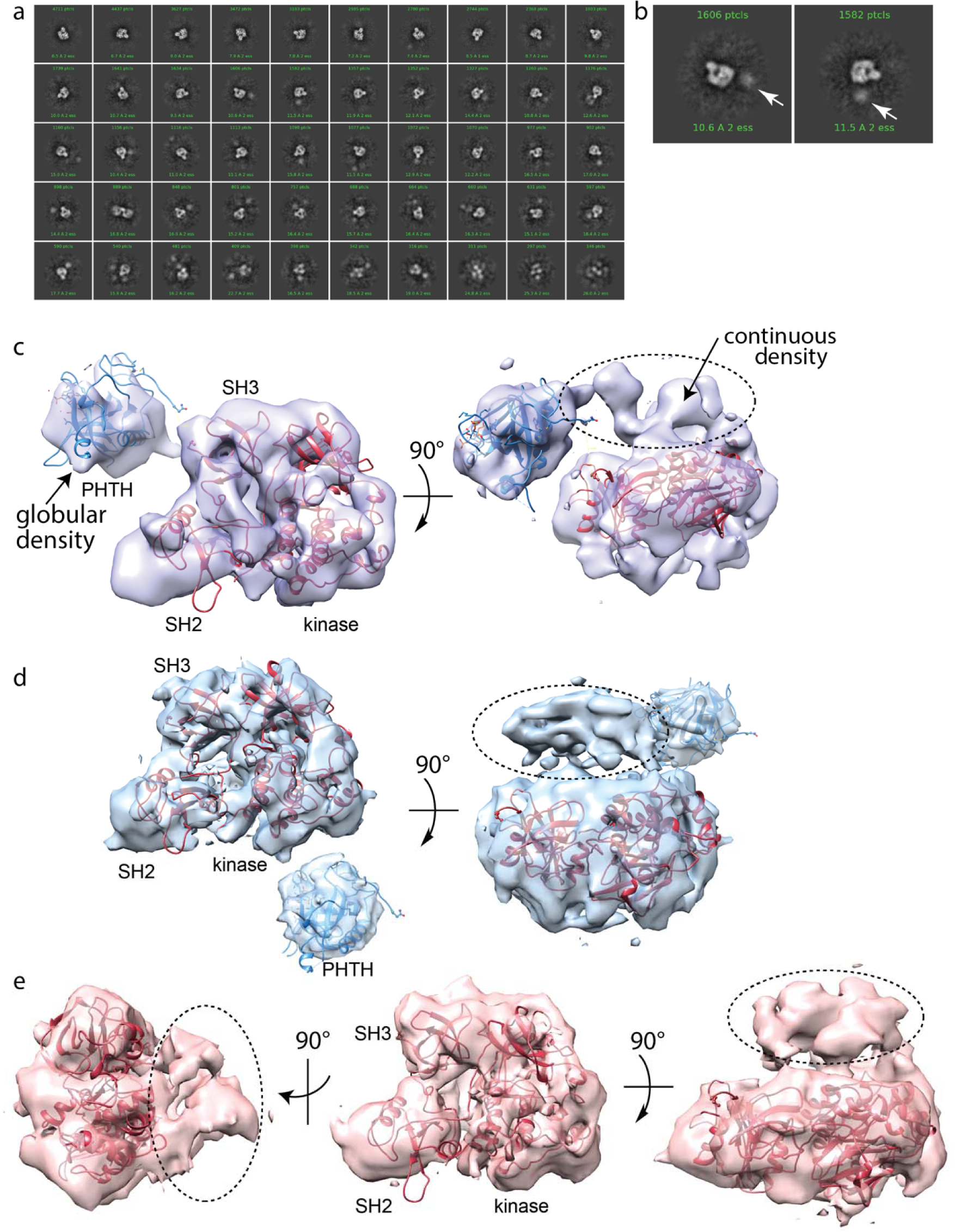
Full-length BTK CryoEM analysis. (a) 2D class averages of full-length BTK. (b) Representative 2D class averages. White arrow indicates extra density adjacent to the BTK SH3-SH2-kinase core density. (c,d,e) Three final 3D reconstructions (see workflow in Fig. 4-Figure supplement 1). (c) Two views of the Class 0 map with the SH3-SH2-kinase fragment (PDB: 8GMB) fit into the larger density and monomeric PHTH domain (PDB: 1B55) fit into the smaller globular density. Continuous density between the large and small density is indicated with a dashed circle. (d) Two views of the Class 1 map with globular density fit as described for (c). The smaller globular density is located in a distinct position with respect to the SH3-SH2-kinase core compared to that shown in (c). Additionally, unmodeled density is observed adjacent to the BTK SH3-SH2-kinase core (dashed circle); the position is similar to the continuous density observed in Class 0. (e) Three views of the Class 3 map with fitted BTK SH3-SH2-kinase core and unmodeled density that is in the same location as that in (d). EMDB accession codes are as follows: EMD-40585, EMD-40586, EMD-40587. Map fitting without user input was also carried out using Situs [71]. The result of that fitting is comparable to results obtained using Chimera.

Two of the three 3D reconstructions (Class 0 and Class 1) show density possibly representing the N-terminal PHTH domain and adjacent PRR linker region (Fig. 4c,d). The PHTH domain structure fits into the extra globular density that sits outside of the SH3-SH2-kinase core, and in one reconstruction, continuous density is observed from the SH3-SH2-kinase core to the globular domain (Fig. 4c). The remote globular density is not visible in Class 3, but similar to Class 0 and 1, continuous unresolved density next to the SH3-SH2-kinase core is present possibly indicating the location of the PRR linker region that is covalently connected to the SH3 domain (Fig. 4e).

Hydrogen deuterium exchange mass spectrometry (HDXMS) data on wild type, full-length BTK (Fig. 4-Figure supplement 2) are consistent with an exposed PRR linker region surrounded by two globular folded domains (PHTH and SH3). Deuterium uptake reaches its maximum at the earliest timepoints for peptides spanning the linker between PHTH and SH3 consistent with high solvent accessibility and the lack of a well-defined hydrogen-bonding network. In contrast, the deuterium uptake curves for peptides derived from all other regions of BTK (PHTH, SH3-SH2, kinase domains) are consistent with folded domains.

## Pursuing the N-terminal PHTH domain

Results from crystallography and cryo-EM presented here suggest the PHTH domain is not stably associated with the autoinhibited SH3-SH2-kinase core of BTK but instead samples a broad conformational ensemble within the context of full-length BTK that may include transient interaction with the autoinhibited core. Indeed, we and others have previously implicated the PHTH domain in autoinhibitory contacts with the kinase domain of BTK [5, 10, 11]. Specifically, we have mapped an interaction between the PHTH domain and the activation loop face of the BTK kinase domain (Fig. 1e) using solution methods. As well, the BTK fragment lacking the PHTH-PRR region is more active than the full-length protein [5] and the same work suggested that mutation of specific surface exposed sidechains within the PHTH domain (R133 and Y134) results in modest activation of BTK. The latter was based on the tightly tethered PHTH-kinase crystal structure (Fig. 1d, PDB: 4Y93) [5]. The location of the PHTH domain in the tightly tethered structure (PDB: 4Y93) is sterically incompatible with the autoinhibited SH3-SH2-kinase structure, and so the authors used molecular dynamics simulations to arrive at a model for full-length autoinhibited BTK that accommodates both the SH3 and PHTH domain on the kinase N-lobe [5]. Given the short linker used in that work, the steric clash between SH3 and PHTH domains in the two separate structures, and the fact that our x-ray and cryo-EM analyses of full-length BTK don’t reveal precise PHTH autoinhibitory contacts, we decided to pursue the role of PHTH in BTK autoinhibition further by crystallizing a redesigned tethered PHTH-kinase construct.

The first step in our redesign was to lengthen the linker between PHTH and kinase domain to reflect the flexibility and distance between the N- and C-terminal domains of BTK. We inserted GGSGG repeats that were either 22 or 32 amino acids in length (Fig 5a). This linker entirely replaced the PRR, SH3, SH2, and SH2-linker sequence. We intentionally excluded the SH2-kinase linker from the PHTH-kinase construct since this linker, in particular L390, mediates autoinhibitory contacts between the SH3 domain and the kinase domain N-lobe. In addition, the new ^1^PHTH^171^-G(GGSGG)_4/6_G-^396^kinase^659^ constructs included PHTH β5-β6 loop mutations (Q91A, I92A, I94A and I95A) to prevent formation of the “Saraste dimer” since that structure is associated with BTK activation [9]. As before, the K430R and ITK activation loop mutations were incorporated to facilitate crystallization. The GDC-0853 inhibitor was included in all crystallization conditions. Both redesigned constructs yielded crystals in the same crystallization conditions (see Materials and Methods), but crystals from the construct containing the 22 amino acid G(GGSGG)_4_G linker yielded larger and higher quality diffracting crystals compared to the construct containing the longer 32 amino acid linker.

**Figure 5.**
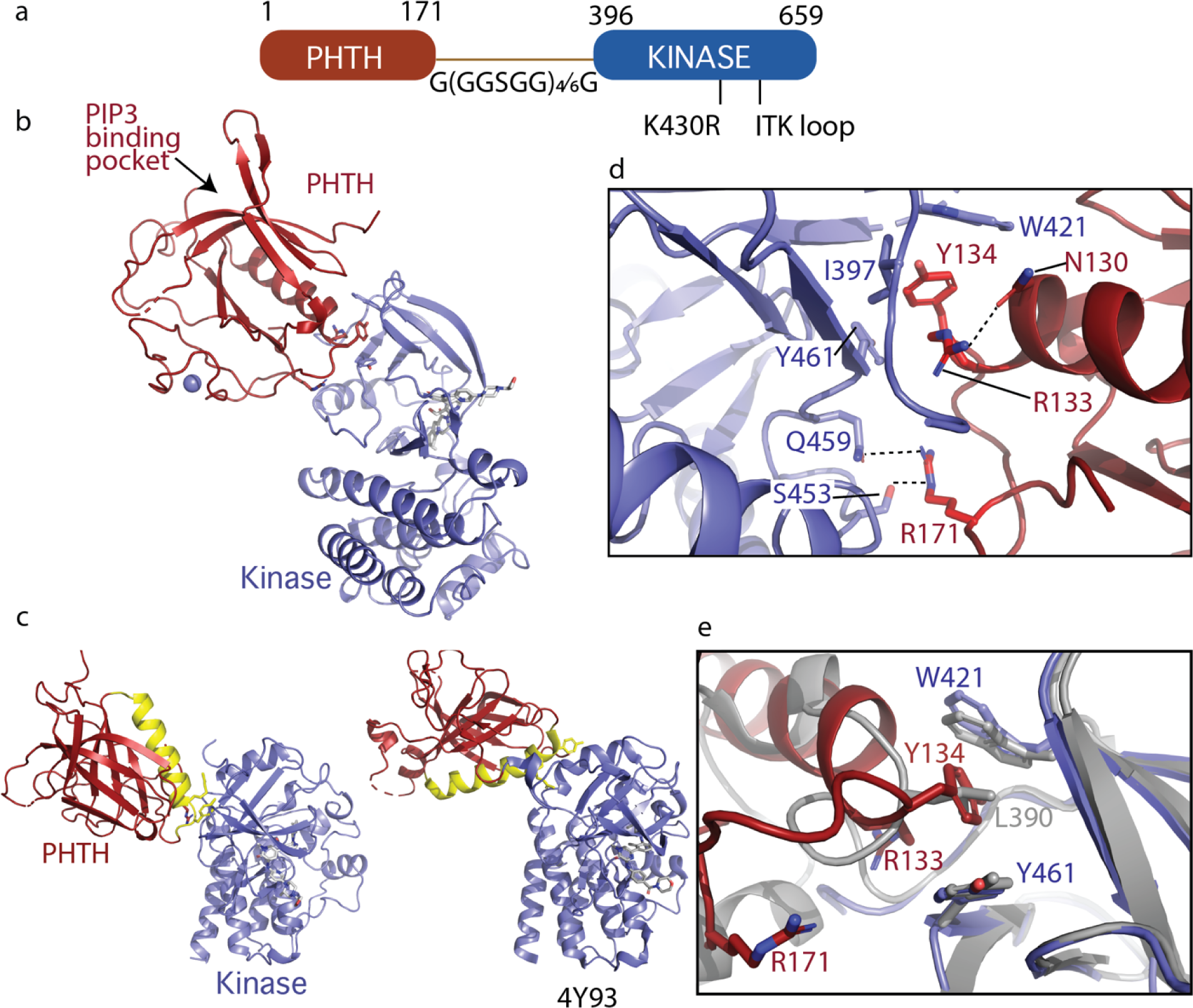
Crystallization of loosely tethered PHTH-kinase. (a) Architecture of the loosely tethered PHTH-kinase constructs used for crystallography. (b) Crystal structure of the PHTH-kinase protein (PDB: 8S93). The PHTH domain (red) docks onto the back of the kinase domain N-lobe (blue). The location of the PIP_3_ binding pocket on PHTH is indicated. (c) Direct comparison of the loosely tethered PHTH-kinase structure solved here (left, PDB: 8S93) and the more tightly tethered PHTH-kinase structure solved previously (right, PDB: 4Y93, right). The PHTH domain helix is colored yellow and the kinase domains are in the same orientation to emphasize the difference between the PHTH domains in the two structures. (d) PHTH/kinase interface. PHTH sidechains R133, Y134 and R171 (red) make contacts to the BTK kinase domain (blue). Dotted lines indicate hydrogen bonds. (e) Close-up view of the hydrophobic stack (flanked by W421 and Y461) on the kinase domain N-lobe. The PHTH Y134 residue (red) inserts into the hydrophobic stack (blue) in the loosely tethered PHTH-kinase structure solved here while L390 from the SH2-kinase linker (gray) completes the hydrophobic stack in the previously solved PHTH-kinase structure (PDB: 4Y93).

The structure of the PHTH-G(GGSGG) _4_G-kinase bound to GDC-0853 was determined to 2.1Å (Fig. 5b). The kinase domain adopts the inactive conformation similar to other structures of inactive BTK (RMSD of 0.43 Å relative to the structure of the isolated inactive kinase domain (PDB: 5VFI) and 0.93 Å relative to the structure of autoinhibited SH3-SH2-kinase (Fig. 3c)). The PIP_3_ binding pocket is sterically accessible (Fig. 5b) as was observed in the tightly tethered BTK PHTH/kinase complex. Otherwise, the orientation and specific contacts between the PHTH and kinase domains in this new, loosely tethered complex (PHTH-G(GGSGG) _4_G-kinase) are quite distinct from the previously solved structure of the PHTH domain tethered to the kinase domain (PDB: 4Y93) (Fig. 5c). In the structure solved here (Fig. 5c, left), the longer linker between domains allows the PHTH domain to bind to the β-strands of the kinase domain N-lobe instead of the C-helix binding site defined previously [5]. Three PHTH domain side chains, R133, Y134 and R171, provide key interactions with the kinase domain in this new structure (Fig. 5d). The Y134 sidechain inserts into the hydrophobic stack [13, 18] created by W421 and Y461 on the kinase domain N-lobe in a manner that mimics the L390 sidechain position in the autoinhibited SH3-SH2-kinase structure (Fig. 5d,e). The guanidinium group of R171 participates in hydrogen bonds with the side chains of S453 and Q459 in the kinase domain (Fig. 5d). The guanidinium group of R133, in contrast, points back into the PHTH domain forming a hydrogen bond with the PHTH N130 side chain (Fig. 5d) while the hydrophobic methylene groups of the R133 side chain contact I397 and Y461. The 22-residue GS linker is only partially visible and is not involved in the PHTH-KD interaction. The total solvent-excluded surface area of the PH-KD interface is 753.8 Å^2^. To further assess the importance of the unique PHTH/kinase interaction captured in this new structure we pursued mutational analysis combined with functional assays.

## Probing the functional role of PHTH/kinase interface

Using a coupled kinase assay, we tested the importance of R131, Y134 and R171 in regulating the activity of BTK. Since the R133/Y134 mutation was also the subject of the previous work examining the regulatory role of the PHTH domain [5], we examined single point mutations (R133E, Y134E and R171E), the double mutant studied previously (R133E/Y134E), and the triple mutation (R133E/Y134E/R171E) in an otherwise wildtype, active version of the PHTH-kinase construct (^1^PHTH^176^-G(GGSGG)_4_G-^384^kinase^659^). BTK kinetics have been extensively characterized previously [19], and our data show the same lag-phase and non-linear time course due to the activating effect of autophosphorylation on Y551 (Fig. 6a,d,g). In an effort to approximate catalytic rates of the mutant and wild type BTK proteins, we fit a line to the data within a short time window for each progress curve (Fig. 6b,e,h) and compare time to a threshold value of ADP produced in the assay (Fig. 6c,f,i). The data for the PHTH-kinase construct show that neither the single mutations nor the double R133E/Y134E mutation alter the rate of the kinase reaction (Fig. 6b) or the time to 3000 pmol ADP (Fig. 6c). In contrast, mutation of all three PHTH domain side chains (R133E/Y134E/R171E) increases BTK activity (Fig. 6a,b) and shortens the time to 3000 pmol ADP (Fig. 6c). These data are consistent with the conclusion that the interface captured in the crystal structure serves an inhibitory role at least in the tethered PHTH-kinase complex.

**Figure 6.**
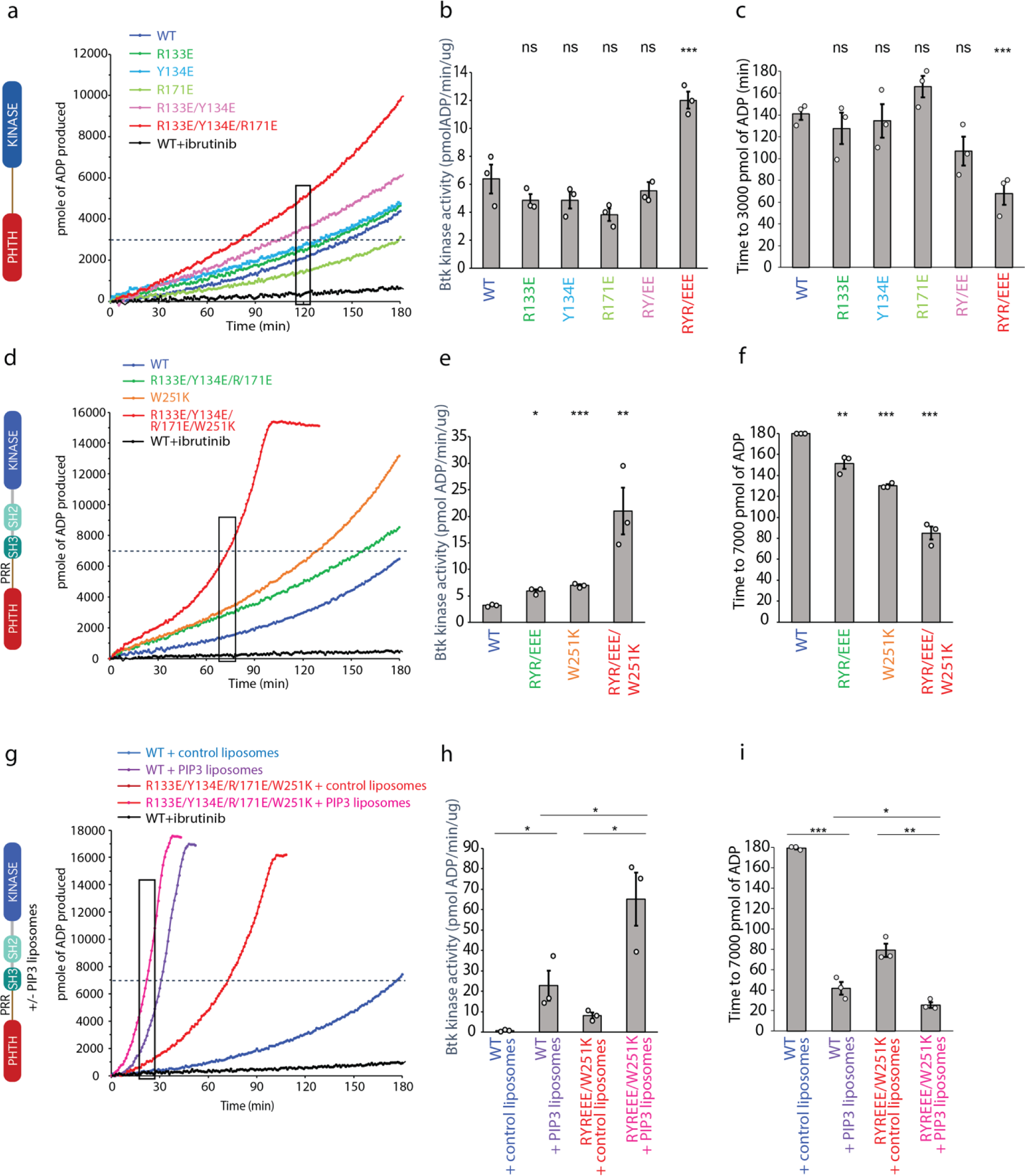
BTK activity assays. (a-c) Representative progress curves, catalytic rate comparisons, and time to threshold ADP for the PH-KD construct. Wild-type BTK PHTH-kinase protein is compared with single, double and triple mutants to probe the PHTH/kinase interface. Ibrutinib inhibition leads to reduction of ADP production (black curve in all experiments). (d-f) Representative progress curves, catalytic rate comparisons, and time to threshold ADP for full-length BTK. Wild type BTK is compared to the following full-length BTK mutants: R133E/Y134E/R171E, W251K, or W251K/R133E/Y134E/R171E. (g-i) Representative progress curves, catalytic rate comparisons, and time to threshold ADP for full-length WT BTK and R133E/Y134E/R171E mutant in the presence of either control or PIP_3_ liposomes. (a,d,g) Representative progress curves of ADP production by BTK are from one of the three independent experiments, and each data point is the average of at least two replicates. (b,e,h) Bar graphs ((b,e,h) represent the average kinase activity rate ± standard error calculated from the boxed region of the corresponding progress curves. Bar graphs (c,f,i) represent the average time to a threshold value of ADP, indicated by dashed line on progress curves. Open circles on all bar graphs represent specific values in each independent experiment. For reactions for which the threshold ADP value is not reached (WT BTK in panels f and i) the values are reported as 180 minutes. The effect of mutations compared with the wild-type BTK was evaluated by Student’s t test (*: P < 0.05; **: P < 0.01; ***: P < 0.001; ns, not significant).

We next tested the same PHTH mutations in the context of full length BTK (Fig. 6d-f). Comparing the activity of wildtype full-length BTK to the R133E/Y134E/R171E mutant, we find that disrupting the PHTH/kinase interface results in a slight increase in the activity of full-length BTK (Fig 6d,e). Since our new PHTH-kinase structures does not contain L390, and instead Y134 fulfills the hydrophobic stack (Fig. 5e), we reasoned that in the full-length protein the PHTH domain may only exert its effect upon release of L390 from the N-lobe (likely concomitant with release of the SH3 domain). We therefore tested the effect of the PHTH mutations in the context of an additional mutation (W251K) that breaks the SH3 contacts with the SH2-linker (contains L390) and kinase domain N-lobe. Mutation of W251K in the SH3 domain of full-length BTK by itself leads to an increase in BTK activity similar to that of the R133E/Y134E/R171E mutation (Fig. 6d,e). Mutation of W251 to lysine in combination with the R133E/Y134E/R171E mutation results in a more active kinase (Fig 6d,e) and shorter time to the threshold ADP level (Fig. 6f) compared to constructs in which the autoinhibitory SH3 domain is intact. These data are consistent with the idea that the PHTH/kinase interaction and the SH3/SH2-linker/N-lobe contacts are mutually exclusive and that both interactions separately serve to allosterically inhibit the catalytic activity of the BTK kinase domain likely via stabilization of the hydrophobic stack.

## Effect of BTK PHTH mutations on activation by phospholipid

PH domains are well characterized phospholipid binding domains and the BTK PHTH domain has been studied extensively in this regard [20]. PIP_3_ binding promotes trans-autophosphorylation, presumably through formation of the PHTH Saraste dimer on the two-dimensional membrane surface. Given the involvement of the PHTH domain in membrane association and dimerization, we tested whether the activating mutation in the PHTH domain (R133E/Y134E/R171E/W251K) has any effect on PIP_3_ mediated activation (Fig. 6g-i). We find that wildtype BTK and the active R133E/Y134E/R171E/W251K BTK mutant are both activated (Fig. 6g,h) and reach threshold ADP levels more rapidly (Fig. 6i) in the presence of PIP_3_ liposomes compared to liposomes that do not contain PIP_3_. Moreover, compared to wild type BTK, the activating mutations in the PHTH domain (R133E/Y134E/R171E) result in increased activity even in the presence of PIP _3_ (Fig 6g,h). Thus, the autoinhibitory effect of the PHTH domain is detected both in the context of non-membrane associated BTK and when BTK is activated by PIP_3_ binding to the PHTH domain.

## BTK kinase dimer structure

The focus of previous work examining BTK autophosphorylation at the PIP_3_ containing membrane surface on has been on the PHTH/PIP_3_ interaction and subsequent PHTH dimerization [20]. PHTH mediated dimerization at the membrane likely promotes a switch from the inactive to active kinase domain conformation followed by trans autophosphorylation of the BTK kinase domain. Consistent with this activation model we have now crystallized the BTK kinase domain in a form that captures a dimer structure where activation loops are swapped to allow for trans autophosphorylation.

The crystallization construct used to capture this new conformation of the BTK kinase domain included mutation of Y551E to mimic the phosphorylated activation loop, mutation of L390G to disfavor autoinhibitory contacts, and dasatinib bound to the active site [21, 22]. The kinase domain forms a face-to-face dimer in the crystal (Fig. 7a) with two intermolecular interfaces; the β3/C-helix loop in the N-lobe forms one interface (Fig. 7b) and the domain swapped activation loop and G-helix mediate an extensive interface between the C-lobes (Fig. 7c). In the N-lobe, the C-helix adopts an intermediate conformation between the previously solved active BTK kinase domain (PDB: 3K54) with the C-helix ‘in’ state and the inactive structure of the BTK kinase domain (PDB: 3GEN) with C-helix ‘out’.

**Figure 7.**
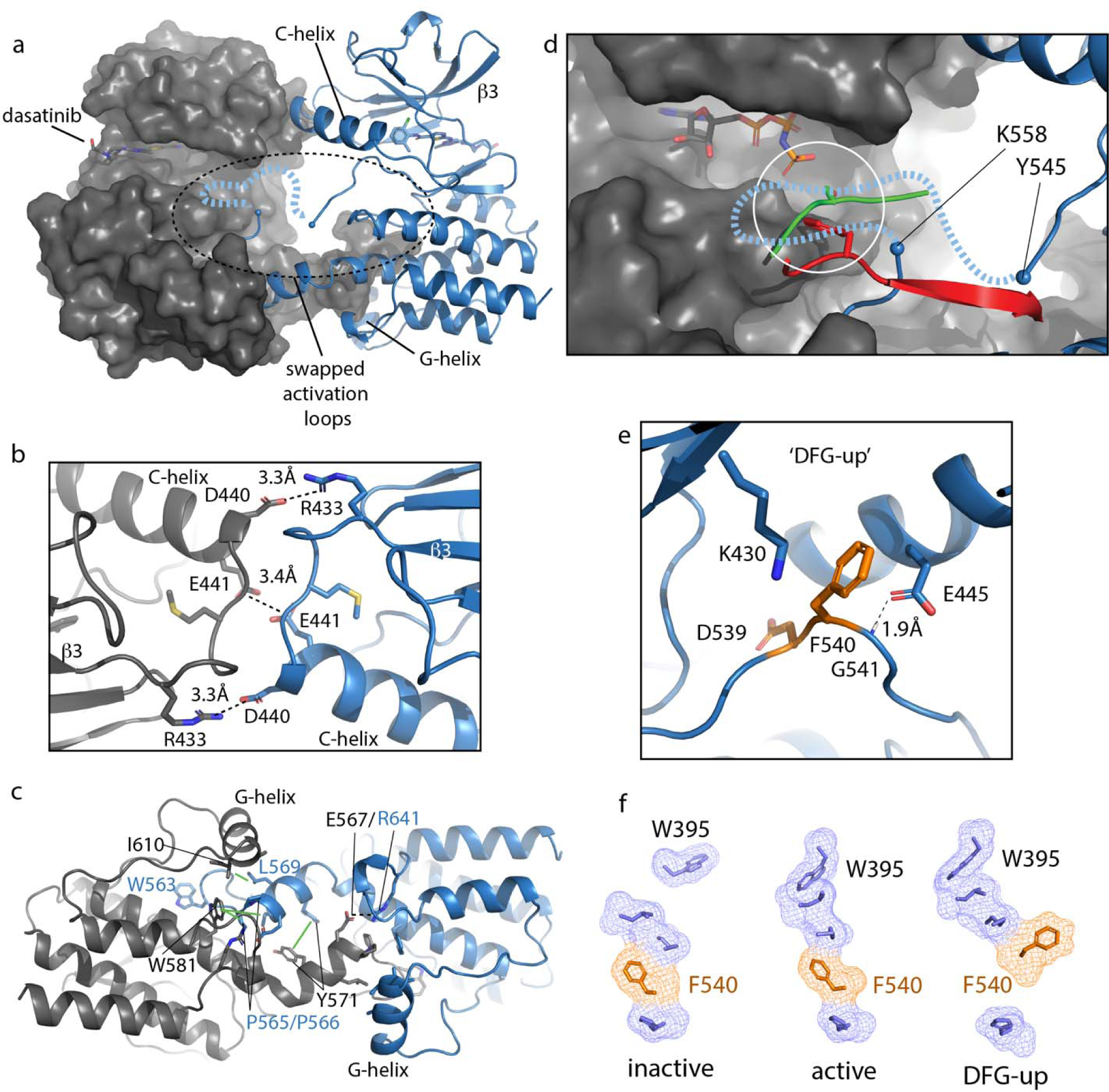
BTK kinase dimer. (a) Crystal structure of the BTK kinase domain dimer (PDB: 8S9F). Monomers are shown in gray and blue with one represented in cartoon and the other surface rendered. The C-helix, β3 strand, G-helix and bound dasatinib are labeled. The region containing the swapped activation loops is indicated with a dashed circle). The portion of the activation loop for which electron density is missing is indicated with a dashed blue line. (b,c) Sidechain interactions mediating the N-lobe and C-lobe dimer interfaces, respectively. (d) Close-up view of activation loop of one monomer extending into the active site of the other monomer. Electron density is absent between Y545 and K558 (indicated with blue spheres). Dashed line indicates possible path for the 13 missing residues that contain Y551. Superimposed on the BTK dimer structure are the PKC kinase domain structure bound to substrate (PDB: 4DC2, green) and insulin-like growth factor 1 receptor kinase bound to substrate (PDB: 1K3A, red); kinase domains are excluded for clarity. The serine and tyrosine phosphoacceptors on these substrates are positioned close to the putative location of BTK Y551 (white circle). (e) Unusual ‘DFG-up’ conformation. In the BTK kinase dimer, F540 inserts between K430 and E445 preventing formation of the salt bridge associated with active kinases. (f) Comparison of regulatory spine structures for active BTK kinase domain (PDB: 3K54), inactive BTK (PDB: 3GEN) and the ‘DFG-up’ structure solved here (PDB: 8S9F). F540 is orange and other R-spine residues are in blue. W395 is at the top of the R-spine in BTK; the ‘DFG-up’ configuration stabilizes the active rotamer of W395 [27, 72].

The activation loop of one monomer in the face-to-face BTK dimer structure extends into the active site of the other monomer but electron density is missing for residues 546 to 559, which includes the Y551 autophosphorylation site (Fig. 7d). Nevertheless, superimposing two different substrate bound kinase structures (tyrosine and serine phospho-acceptors) onto the BTK kinase domain (Fig. 7d) shows that Y551 from one BTK kinase domain monomer would easily reach a viable phosphor-acceptor position in the active site of the opposing monomer. The swapped BTK dimer structure may therefore represent a conformation along the trans-autophosphorylation activation pathway [23].

The canonical Lys/Glu salt bridge with active kinase domains does not form in the BTK kinase domain dimer due to the unusual conformation of the phenylalanine side of the DFG motif (F540) (Fig. 7e). The F540 sidechain intercalates between the Lys/Glu sidechains adopting what has previously been termed ‘DFG-up’ [24, 25] or ‘DFGinter’ [26]. The C-helix is therefore held in an intermediate state between the C-helix in and C-helix out conformations associated with active and inactive kinases, respectively. In addition to the unusual position of F540, D539 of the DFG motif points inward, orientated approximately 180° opposite from that in the DFG-in conformation of the active kinase and E445 on the C-helix forms a hydrogen bond with the backbone amide of G541 in the DFG motif instead of forming a salt bridge with K430 (Fig. 7e). It is also interesting to compare the regulatory spine (R-spine) for this BTK kinase dimer and the R-spine in previously solved active and inactive structures of BTK (Fig. 7f). The active BTK structure shows close packing between the top of the R-spine and W395, which adopts the rotamer conformation required for active BTK [27]. Inactive BTK, in contrast, shows no contact between W395 and the R-spine (Fig. 7f); interestingly, the DFG conformation is quite similar between active and inactive BTK. The ‘DFG-up’ configuration in the kinase domain dimer structure is accompanied by the W395 rotamer that is associated with active BTK (Fig. 7f). That said, it is also possible that the DFG status is highly dependent on the nature of the bound drug. In fact, a previous structure of another TEC family kinase, BMX, reveals the same DFG-up state and the kinase domain in that case is also bound to dasatinib [28].

## Discussion

The findings reported here provide new insights into autoinhibition and allosteric control of the full-length, multi-domain BTK protein. The data advance our understanding of BTK regulation as many previous studies did not interrogate the structure of the entire full-length protein. In pursuing full-length BTK, we find that missing electron density in crystals of full-length BTK, elongated SAXS envelopes, and visualization of the PHTH domain by cryoEM support a model where the N-terminal BTK PHTH domain populates a conformational ensemble surrounding the compact/autoinhibited SH3-SH2-kinase core (Fig. 8a). In previous work probing allosteric regulation of full-length BTK [6], we consistently find that perturbations to the conformational state of the SH3-SH2-kinase core have no effect on the hydrogen/deuterium exchange behavior of the PHTH domain consistent with the dynamically independent PHTH domain observed here. Contrary to models of autoinhibited multidomain proteins where each globular domain is sequestered into a compact core, the autoinhibited state of full-length BTK might instead feature a conformationally heterogeneous PH domain that is connected to a compact core through a flexible linker and likely contacts the kinase domain only transiently. Similar conformational heterogeneity has been reported for the PH domains of phospholipases C ε and β [29].

**Figure 8.**
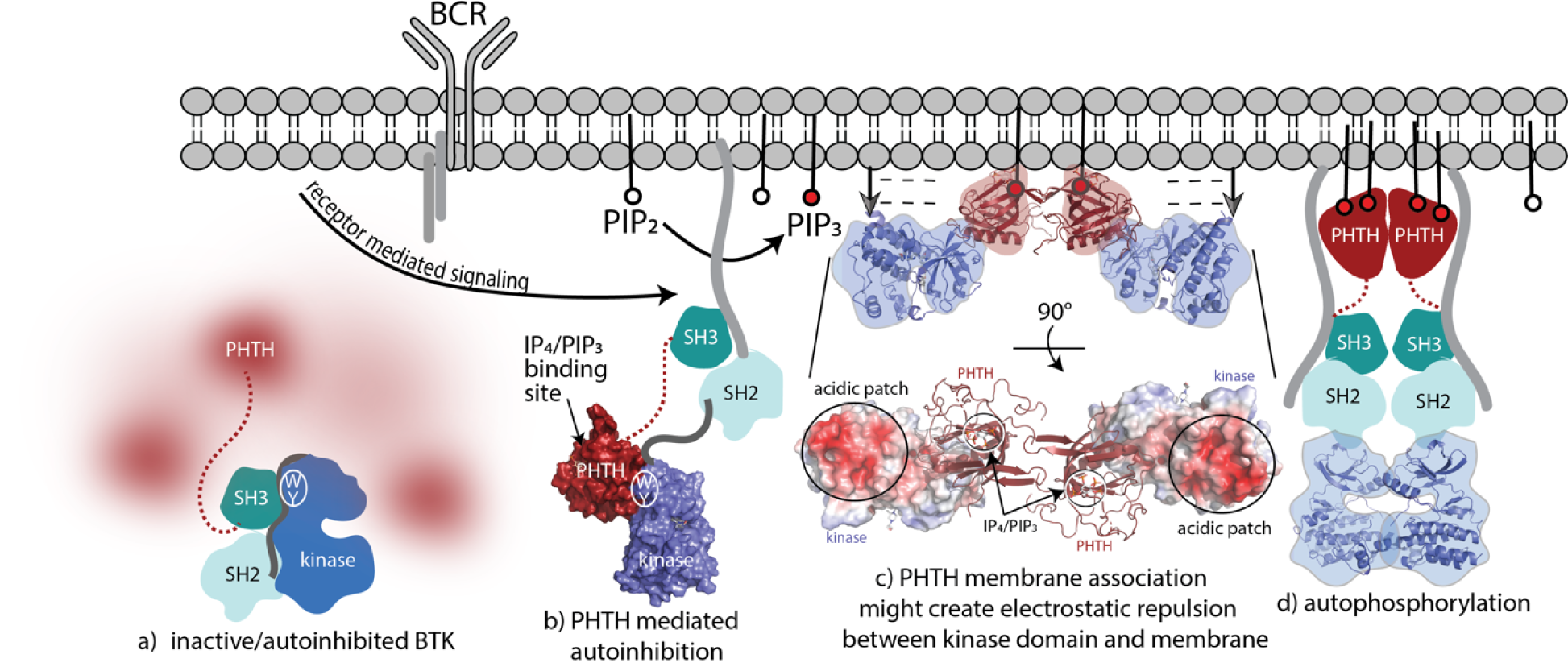
BTK activation model. (a) Inactive, autoinhibited BTK, the conformational heterogeneity of the PHTH domain is indicated in fuzzy red. The hydrophobic stack residues, W421 and Y461, are indicated on the kinase domain N-lobe as [math]. (b) Engagement of the BTK SH3 and SH2 domains with exogenous ligands would allow for the PHTH domain to adopt its autoinhibitory pose. Surface rendering of structure solved here (PDB: 8S93) is included, accessibility of the IP_4_/PIP_3_ binding site is shown, and hydrophobic stack resides are indicated as in (a). (c) Two PHTH-kinase structures are superimposed on the Saraste PHTH dimer (PDB: 1B55). The top model indicates potential for electrostatic repulsion between membrane and BTK kinase domain in this arrangement (negative charges indicated by and arrows suggest unfavorable interactions between negatively charged surfaces). A 90° rotation (bottom) shows the acidic patch on the kinase domain and the PIP_3_ binding sites on the PHTH domain (circled) are on the same surface of the structure. (d) Release of all autoinhibitory contacts and dimerization of the BTK kinase domain (PDB: 8S9F) lead to autophosphorylation on Y551 in the activation loop of each kinase domain.

Another PH domain containing kinase, AKT1, has been studied extensively [30-35] and many of the structures that have been solved to elucidate autoregulation by the PH domain are seemingly biased by allosteric inhibitors. A recent crystal structure [32] provides a more unbiased view of the potential autoinhibitory interface between the PH and kinase domains of AKT; the PH domain binds to the activation loop face of the kinase domain in a manner that is quite similar to our earlier solution studies of both ITK and BTK (see Fig. 1e) [6, 10, 11]. Like AKT, the BTK PH/kinase interface detected in solution studies masks the PIP_3_ binding site on the PH domain. However, consideration of all data to date suggests that regulatory interactions between the BTK PHTH and kinase domains are transient. Previously characterized contacts between the PHTH domain and the activation loop face [11] or between the PHTH domain and the kinase domain N-lobe [5] may all exist within the conformational ensemble of a range PH/kinase domain interactions.

Non-PH domain containing kinases present similarly dynamic N-terminal regions that exert regulatory control over catalytic activity. The N-terminal region of the SRC kinases has been invoked in ‘fuzzy’ intramolecular regulatory interactions with the Src module (SH3-SH2-kinase) core [36]. For SRC it has been suggested that the intrinsically disordered N-terminal regions increase the flexibility of the SH3-SH2-kinase core to promote ligand binding interactions [37]. The N-terminal “cap” region of ABL is another example of a long, disordered N-terminal region; myristoylation results in a key autoinhibitory contact between the N-terminal myristate and the ABL kinase domain C-lobe [38-41]. Within BTK, we have previously shown that the proline-rich region (PRR) in the linker between the PHTH and SH3 domains competes with the SH2-kinase linker for binding to the SH3 domain thereby shifting the conformational equilibrium of the autoinhibited core to a more open configuration [6, 42]. Indeed, BTK SH3 displacement from the autoinhibited core via interaction with the PRR may be a prerequisite for the PHTH domain to take up the autoinhibitory pose on the kinase domain N-lobe defined here (Fig. 8b). The extent to which the intrinsically disordered PRR region of BTK participates in additional, perhaps fuzzy, interactions with the BTK SH3-SH2-kinase region remains to be investigated. As well, interactions of the N-terminal region of BTK with both positive and negative regulators [43-45] may alter the dynamic characteristics observed here. It is also possible that the dynamic linker between the PHTH and SH3 domains of BTK may mediate phase separation [46] and clustering into signaling competent condensates. While these questions need further study, it is notable that the SRC and TEC kinases seem to share dynamic regulatory features in their N-termini despite drastically different primary structures.

Transition from fully autoinhibited to active BTK following receptor activation can be considered in light of the structural insights now available (Fig. 8). The BTK hydrophobic stack residues, W421 and Y461, are a binding site for both the L390 sidechain on the SH2-kinase linker [13] (Fig. 8a) and Y134 on the BTK PHTH domain (Fig. 8b). Both interactions allosterically inhibit BTK catalytic activity by stabilizing the inactive kinase domain conformation. The SH3/L390/hydrophobic stack interaction is present in the autoinhibited Src module and release of the regulatory SH3/SH2 domains opens the kinase N-lobe for secondary inhibitory interactions with the PHTH domain (Fig. 8b). Once PIP _3_ is made available by the action of PI3K, the PHTH binds to PIP _3_ in the membrane and likely dimerizes through the Saraste dimer. To explore possible consequences of BTK dimerization at the membrane, we superimposed the PHTH-kinase structure onto the PHTH Saraste dimer (Fig. 8c). Viewing the resulting dimer arrangement from the inner membrane surface shows a large acidic patch on the BTK kinase domains parallel to the bound IP _4_ ligands. If the PHTH-kinase complex remains intact upon membrane association (Fig. 8c), electrostatic repulsion between the negatively charged BTK kinase domain and the negatively charged membrane surface might result in dissociation of the inhibitory BTK PHTH/kinase interaction and subsequent dimerization-mediated autophosphorylation in the kinase domain (Fig. 8d). Indeed, mutation of the PHTH domain residues that contact the kinase domain led to more rapid PIP _3_ mediated activation of full-length BTK (Fig. 6g-i) suggesting the PHTH autoinhibitory structure interaction might remain intact as BTK associates with PIP _3_. This is consistent with a recent report that Grb2 association with the PRR of BTK results in higher levels of BTK activity at the PIP _3_ membrane possibly by destabilizing autoinhibited BTK [47].

The autoinhibitory role of the PHTH domain in maintaining the BTK kinase domain in its inactive state until assembly at the membrane provides new insight into oncogenic forms of BTK. Specifically, the oncogenic p65BTK isoform lacks most of the PHTH domain and is expressed in colorectal carcinoma and some glioblastoma patients [48]. As well, an extended transcript yields BTK-C, an isoform that contains additional residues at the N-terminus and is expressed in breast and prostate cancer cells and [49, 50]. It is not fully understood to what extent and through what mechanisms these deletions and extensions of the PHTH domain affect BTK regulation [51]. In addition to effects on PIP _3_ binding, and/or interactions with other regulatory proteins, it is also possible that the altered PHTH domain sequences may contribute to oncogenicity by disrupting the autoinhibitory function of the PHTH domain.

The TEC kinases provide a rich tapestry on which multidomain protein regulation and allostery can be studied. The observations here support a model where the BTK PHTH domain visits multiple conformational states and likely rapidly interconverts between states (both detached and associated with the SH3-SH2-kinase core). A model where the primary autoinhibitory contacts are housed within the SH3-SH2-kinase core is consistent with the fact that the SRC kinases lack the PH domain entirely and rely only on their SH3 and SH2 domains for autoinhibition [52]. Despite the temptation to assign a stable autoinhibitory configuration to the BTK PHTH domain, the inherent conformational heterogeneity of the BTK PHTH domain instead suggests the possibility of multifaceted regulatory roles for this single domain. It will be interesting to compare the TEC kinases in this regard. The length of the PRR linker between PHTH and SH3 domain varies across the family, the loops that mediate formation of the BTK PHTH Saraste dimer are absent in the PHTH domains of other TEC kinases, and one TEC kinase, RLK/TXK, lacks the PHTH domain and instead contains an N-terminal region nearly identical in length to that of the SRC kinases. Sub-family specific differences in structure, dynamics, autoregulation, activation, and allostery are likely important for control of distinct signaling pathways in distinct cell types.

## Materials and Methods

### Cloning and constructs

Cloning and mutagenesis in this work were carried by Polymerase Chain Reaction (PCR) with Phusion Hot Start II DNA Polymerase (Thermo Fisher Scientific). For initial protein characterization, full-length (FL) BTK (residues 1-659) was cloned into pET20b vector with a C-terminal 6xHistidine tag (FL BTK-C6H) [6]. For crystallization and activity assays, a N-terminal 6x Histidine-SUMO-tag FL BTK (N6H-SUMO-FL BTK) construct was created by inserting a N-terminal 6x Histidine-SUMO-tag into pET20b FL BTK-C6H by Seamless Ligation Cloning Extract (SLiCE) cloning and removing the C-terminal 6xHistidine by PCR (pET-His-SUMO provided by Eric Underbakke). The ^1^PHTH^171^-G(GGSGG)_4/6_G-^396^kinase^659^ constructs were cloned into pET28b vector with a N-terminal 6x Histidine-SUMO-tag. The BTK kinase domain (residues 382-659) construct (N6H-SUMO KD) was created by deleting residues 1-381 from the pET20b N6H-SUMO-FL BTK plasmid by PCR. All BTK constructs carry the solubilizing Y617P mutation [6]. All constructs created in this work were verified by sequencing at the Iowa State University DNA synthesis and sequencing facility.

### Protein expression and purification

The plasmids were transformed to BL21 (DE3) cells (MilliporeSigma). To express FL BTK-C6H, N6H-SUMO-FL BTK, and N6H-SUMO KD, cells were grown in LB broth (Fisher Scientific) supplemented with 100 μg/mL ampicillin (Fisher Scientific). For the expression of ^1^PHTH^171^-G(GGSGG)_4_G-^396^kinase^659^ and ^1^PHTH^176^-G(GGSGG)_4_G-^384^kinase^659^ fusion proteins (PHTH-KD), cells were grown in LB broth supplemented with 50 μg/mL kanamycin (MilliporeSigma). For expression of any kinase active BTK, pCDFDuet YopH was co-transformed with the BTK plasmids additionally supplemented with 50 μg/mL streptomycin. All BTK constructs were expressed similarly: cells were grown at 37°C until the optical density at 600 nm reached ∼0.6. Protein expression was induced with 1 mM isopropyl β-d-1-thiogalactopyranoside (IPTG) for 24 hours at 18°C. Cells were harvested by centrifugation at 3,000 × g for 20 min, resuspended in lysis buffer (50 mM Tris pH 8.0, 500 mM NaCl and 10% glycerol), and stored at -80°C until use. Cells were thawed with the addition of 0.5mg/ml lysozyme and 1500U of DNAse I (MilliporeSigma) and lysed by sonication on ice at 50% amplitude for 2 min with 0.5-second on and 1.5-second off intervals. The lysate was clarified by centrifugation at 20,000 × g for 1 hour at 4°C, and the supernatant was applied to a Ni-NTA column (QIAGEN) equilibrated with lysis buffer. The column was washed with 20 column volume of lysis buffer with 10 mM imidazole. For FL BTK-C6H, the proteins eluted with lysis buffer with 300 mM imidazole. For N6H-SUMO-tagged BTK constructs, the BTK-bound Ni-NTA resins in disposable columns were capped, resuspended with the lysis buffer with the addition of 5mM TCEP and 1mg of Ulp1 (provided by Eric Underbakke), and incubated O/N. The tagless BTK proteins were obtained by collecting the flow-through from the column and eluting with additional 10mL of lysis buffer. The BTK sample was concentrated, and further purified with HiLoad 26/600 Superdex 200 pg (Cytiva) with SEC buffer (20 mM Tris pH 8.0, 150 mM NaCl, 10% glycerol and 1mM DTT). The purity of the samples was verified by SDS-PAGE. If YopH is still present in the BTK fractions, proteins were further purified with a Source 15Q column using ion-change buffers (Buffer A: 20 mM Tris pH 8.0, 1 DTT, and 50mM; Buffer B: 20 mM Tris pH 8.0, 1 DTT, and 1M NaCl). Purified proteins were concentrated to 5 to 20 mg/mL, flash-frozen in 100ul aliquots with liquid nitrogen, and stored at -80°C.

## Crystallization, structure determination and refinement

### Crystal structure of FL BTK

The FL BTK construct (residues 1 to 659) used for initial crystallization trials contains multiple mutations in different regions of BTK: 1. PHTH domain β5-6 loop mutations: Q91A, I92A, I94A and I95A; 2. PPR region mutations: P189A, P192A, P203A and P206A; 3. SH2-kinase linker mutations: A384P, S386P, T387P, A388P and L390F; 4. Activation loop mutations: L542M S543T, V555T, R562K, S564A, P565S; 5. Kinase inactive mutation: K430R. Tiny needle-shaped crystals of FL BTK were initially obtained from sitting-drop vapor diffusion method by mixing purified FL BTK proteins at 15-20 mg/mL with equal amount of the precipitant solution 15-20% polyethylene glycol (PEG) 3350, 0.1M Bis-Tris Propane, pH7.5, and 0.2M potassium thiocyanate (KSCN) at 4 °C. Larger FL BTK crystals (0.2-0.4mm) were obtained after E298A, K300A, E301A mutations were introduced to the construct based on results from Surface Entropy Reduction prediction (SERp) server [17]. Final FL BTK construct for crystallization (FL BTKsc) contains only the SH2-KD linker mutations (A384P, S386P, T387P, A388P, L390F), activation loop mutations (L542M S543T, V555T, R562K, S564A, P565S; also included in previous solution studies mapping the PHTH/kinase interaction [11]); active kinase salt bridge mutation (K430R) and the SERp mutations (E298A, K300A, E301A). In an effort to reduce the flexibility of the PRR containing linker, five additional full-length BTK constructs with the following deletions: Δ185-194, Δ 185-206, Δ181-206, Δ175-210, and Δ 173-215, were also subject to crystallization and structural analysis, but no electron density was observed for the PHTH domain from any of these crystals.

For x-ray diffraction data collection, crystals were transferred, with six sequential steps with increasing amount of glycerol, to a solution containing 17% PEG 3350, 0.1M Bis-Tris Propane, pH7.5, 0.2M KSCN, and 20% glycerol, and then flash-frozen with liquid nitrogen. X-ray diffraction data were collected at the Advanced Photon Source (APS) beamline 23-ID-B GM/CA and 24-ID-E NE-CAT. Data sets were indexed, merged, and scaled using autoPROC (Global Phasing) [53, 54]. The structure was solved by molecular replacement (MR) using Phaser in the CCP4i suite [55, 56] using the previous SH3-SH2-KD structure (PDB code 4XI2) as the search model. The structures were refined using Phenix [57] and built in Coot [58]. Complete X-ray collection and refinement statistics are provided in Table 1. The figures are prepared with Pymol [59].

**Table 1.**
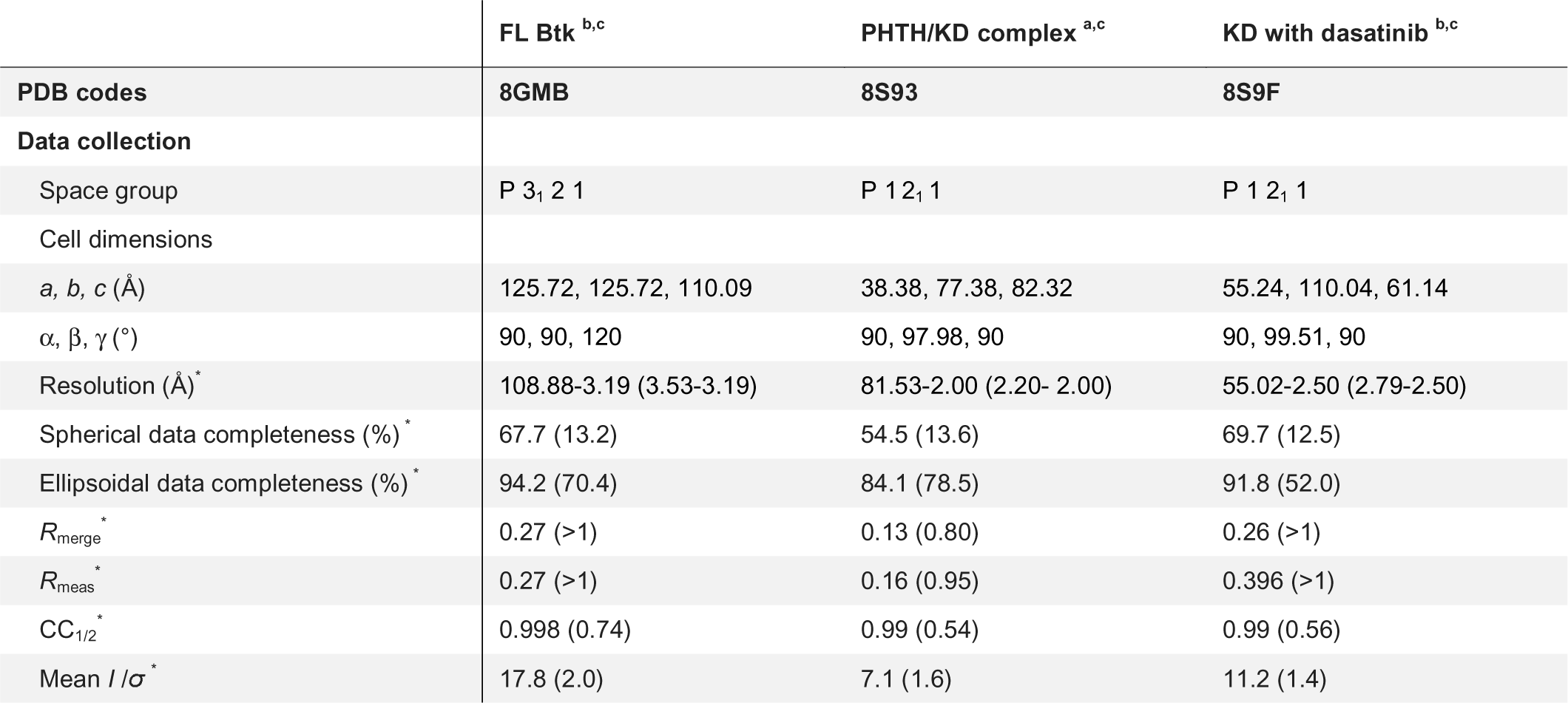
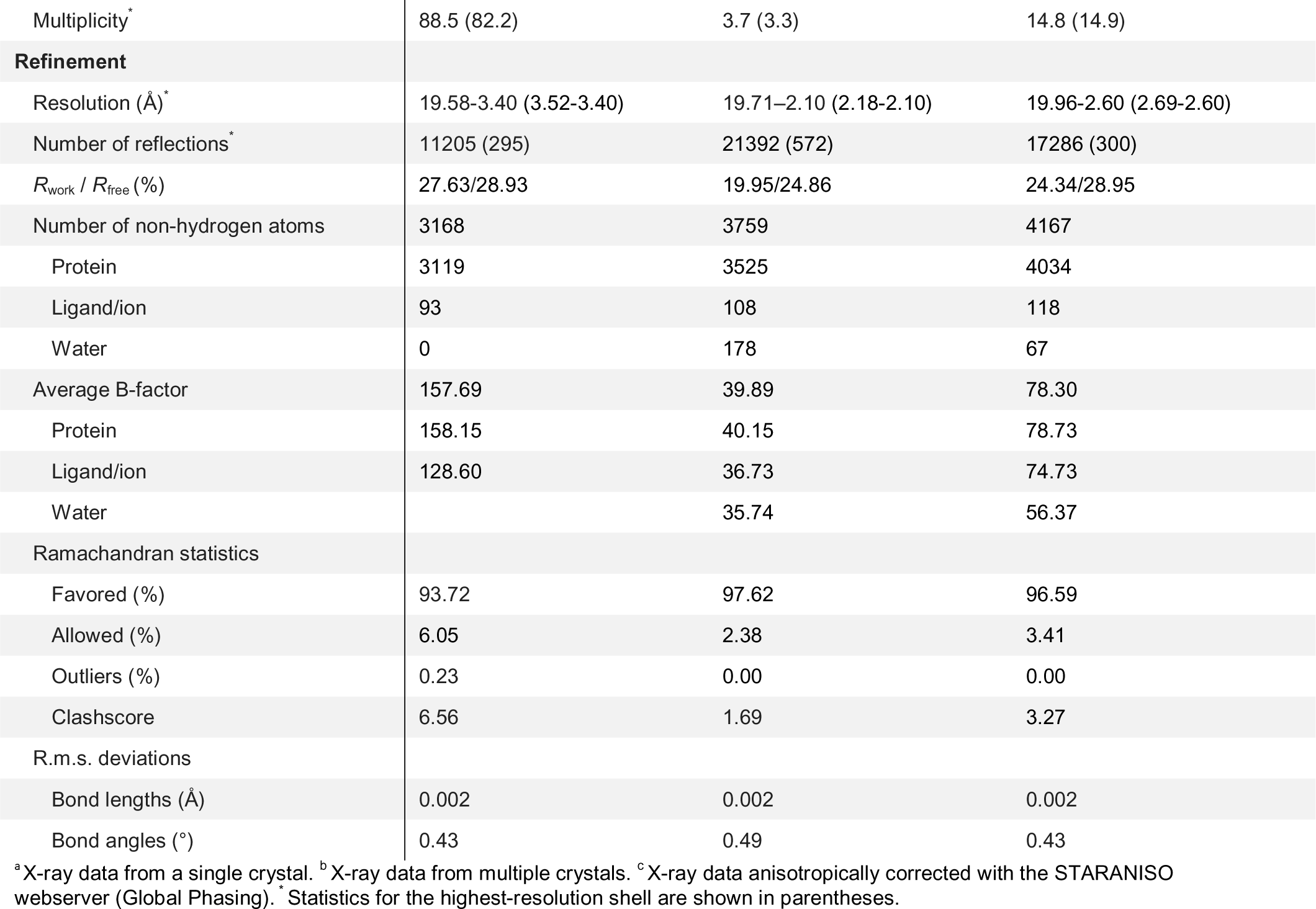
Data collection and refinement statistics.

### Crystal structure of the PHTH-KD complex

PHTH-KD proteins at 10mg/ml were crystalized with hanging-drop method by mixing at 1:1 ratio with precipitant solution, 20% PEG3350, 0.1M Bis-Tris, pH 5.6, 0.2M MgCl _2_. Thin plate-like crystal clusters were harvested and flash-frozen in the cryo-protectant solution (precipitant solution+20% glycerol). X-ray diffraction data were collected at the APS beamline 23-ID-B GM/CA. PH-KD structure was solved by MR with the isolated structures of the PHTH domain (PDB code 1BTK) and the kinase domain bound to GDC-0853 (PDB code 5vfi) [9, 60]. The structures were refined using Phenix [57] and built in Coot [58].

### Crystal structure of the kinase domain bound to dasatinib

BTK KD Y551E/L390G (residue 382-659) at 20mg/ml was crystalized with the sitting-drop method by mixing at 1:1 ratio with precipitant solution, 42% PEG 200, 0.1M HEPES, pH 7.5. Crystals were harvested and flash-frozen in the cryo-protectant solution (precipitant solution+20% glycerol). X-ray diffraction data were collected at the APS beamline 23-ID-B GM/CA and 24-ID-E NE-CAT. The structure of BTK KD bound to dasatinib was solved by molecular replacement with the previous dasatinib-bound KD structure (PDB code 3K54) [22]. The structures were refined using Phenix [57] and built in Coot [58].

All structures were deposited to the Protein Data Bank with accession codes as follows: FL BTK: 8GMB; PHTH-KD: 8S93; BTK KD/dasatinib: 8S9F.

### CryoEM sample preparation, data collection and processing

An aliquot of purified FL BTK (18 mg/ml) was thawed and re-purified with a Superdex 200 10/300 GL column. The peak fractions were either used directly for cryoEM grid preparation or flash-frozen with liquid nitrogen. 3ul of FL BTK at 0.4-0.6 mg/ml were applied to freshly glow-discharged 300-mesh Quantifoil R1.2/1.3 copper grids (EMS), blotted for 3 seconds with the blot force of 3, and plunged into liquid ethane using FEI Mark IV Vitrobot operated at 4°C and 100% humidity. Dataset was collected on a 200 kV Glacios Cryo-TEM electron microscope (Thermo Fisher Scientific) equipped with a Gatan K3 direct electron detector in super resolution mode using Serial EM at a pixel size of 0.45315 Å with total dose of 59 electrons·Å ^-2^ distributed over 50 frames. 755 movies were selected for data processing. Movies were binned 2x to 0.9063 Å during alignment using Patch Motion Correction. CTF was estimated using Patch CTF estimation and particles were picked by blob picker in Cryosparc-Live. All further steps were done in Cryosparc v4.1.2 [61]. Picked particles were extracted with a box size of 386 pix and fourier cropped to a box size of 184 pix. Extracted particles were subjected to multiple rounds of 2D classification resulting in 68,788 particles which were classified into four Ab-initio reconstructions and further refined using non-uniform refinement [62]. All maps were visualized and analyzed using UCSF Chimera [63]. Electron Microscopy Data Bank (EMDB) accession codes: EMD-40585, EMD-40586, EMD-40587

### Hydrogen deuterium exchange mass spectrometry

Prior to Deuterium labeling, wild type, full-length BTK (20 μM) was thawed on ice and then incubated at 23 °C (room temperature) for 1 hour, diluted to 8 μM with buffer (20 mM Tris pH 8.0, 150 mM NaCl, 10% glycerol) and then returned to ice in preparation for deuterium labeling. Deuterium labeling was initiated with an 18-fold dilution (36 μL) into D_2_O buffer (20 mM Tris pD 8.0, 150 mM NaCl, 10% glycerol, 99.9% D _2_O). After each labeling time (10 seconds, 1 minute, 10 minutes, 1 hour, 4 hours) at room temperature, the labeling reaction was quenched with the addition of an equal volume (38 μL) of ice-cold quenching buffer [150 mM potassium phosphate, pH 2.4, H_2_O] and analyzed immediately.

Deuterated and control samples were digested online at 15 °C using an Enzymate pepsin column. The cooling chamber of the HDX system, which housed all the chromatographic elements, was held at 0.0 ± 0.1 °C for the entire time of the measurements. Peptides were trapped and desalted on a VanGuard Pre-Column trap [2.1 mm × 5 mm, ACQUITY UPLC BEH C18, 1.7 μm] for 3 minutes at 100 μL/min. Peptides were eluted from the trap using a 5%-35% gradient of acetonitrile with 0.1% formic acid over 6 minutes at a flow rate of 100 μL/min, and separated using an ACQUITY UPLC HSS T3, 1.8 μm, 1.0 mm × 50 mm column. Mass spectra were acquired using a Waters Synapt G2-Si HDMS ^E^ mass spectrometer in ion mobility mode. The error of determining the average deuterium incorporation for each peptide was at or below ± 0.25 Da, based on deuterated peptide standards.

Peptides were identified from replicate HDMS ^E^ analyses (as detailed in the Supplemental Data file) of undeuterated control samples using PLGS 3.0.3 (Waters Corporation). The peptides identified in PLGS were then filtered in DynamX 3.0 (Waters Corporation) implementing a minimum products per amino acid cut-off of 0.25, at least 2 consecutive product ions (see Supplemental Data file). Those peptides meeting the filtering criteria to this point were further processed by DynamX 3.0 (Waters Corporation). The relative amount of deuterium in each peptide was determined by the software by subtracting the centroid mass of the undeuterated form of each peptide from the deuterated form, at each time point. Deuterium levels were not corrected for back exchange and thus reported as relative [64].

All deuterium uptake values are provided in the Supplemental Data file along with an HDX data summary and comprehensive list of experimental parameters in the recommended [65] tabular format. All HDX MS data have been deposited to the ProteomeXchange Consortium via the PRIDE partner repository [66] with the dataset identifier PXD041657.

### Thermal shift assay

Purified proteins at 0.5mg/mL were mixed with SYPRO™ Protein Gel Stains (Thermo Fisher Scientific) at 1:500 dilution. 20 uL of the protein/SYPRO™ mixture were aliquoted to MicroAmp Fast Optical 96-Well Reaction Plate (Thermo Fisher Scientific). The experiments with temperature ramp from 25-95°C and 0.5°C/step were measured with The Applied Biosystems StepOnePlus™ 96-well qPCR system (Thermo Fisher Scientific). The melt curve data were analyzed with the StepOne Software (Thermo Fisher Scientific) and the melting temperatures of each construct was determined by taking the lowest point of the negative derivative of normalized fluorescence. Three measurements from each mutant were normalized, averaged and plotted with Excel.

### Preparation of liposomes

1,2-dioleoyl-sn-glycero-3-phosphocholine (DOPC), 1,2-dioleoyl-sn-glycero-3-phospho-L-serine (DOPS) and 1,2-dioleoyl-sn-glycero-3-[phosphoinositol-3,4,5-trisphosphate] (tetra-ammonium salt) (PIP_3_) (Avanti Polar Lipids) were dissolved in chloroform at 10mg/ml and stored at -20°C. To prepare control and PIP _3_ liposomes, DOPC, DOPS and PIP _3_ were mixed in glass test tubes at the molar ratio of 80:20:0 and 75:20:5 (DOPC: DOPS: PIP _3_) respectively. Chloroform was removed by blowing a gentle stream of nitrogen gas until no solvent is visible. The lipids were transferred to a vacuum desiccator and dried overnight at room temperature. Dry lipid films were hydrated in a buffer containing 50 mM HEPES pH 7.4, 150 mM NaCl, and 5% glycerol to a final concentration of 12.5 mM. The hydrated liposomes were subjected to three freeze-thaw cycles using liquid nitrogen, followed by 10 passes through 100-nm filters (MilliporeSigma).

### NADH-coupled kinase assay

The kinase activity of BTK was determined using Pyruvate kinase (PK)/lactate dehydrogenase (LDH) coupled kinase assay [67]. 5μl of BTK at 20μM or 5μl of SEC buffer (for blanks) was added to 20 μl of the reaction buffer containing 50 mM HEPES, pH 7.4, 150 mM NaCl, 5% glycerol, 1mM DTT and 1.6 μL of PK/LDH enzymes from rabbit muscle (MilliporeSigma). For the liposome experiments, 200 μM liposomes were added to the both control and kinase reactions. Experiments were performed in triplicates in black 96-well plates and were measured alongside three control reactions for nonenzymatic production of ADP (no kinase added). The reactions were initiated by adding 25 μl of 2xATP/NADH buffer, containing 1mM NADH, 2mM PEP, 2mM ATP and 20mM MgCl_2_ in reaction buffer and measured using the Synergy HT Microplate Reader (BioTek) by monitoring the fluorescence of NADH at 460 nm at 22°C with 1-min intervals for 3 hours. The absolute concentration of NADH stock at ∼40mM was determined by measuring its absorbance at 340 nm (ε340 = 6220 M-1cm-1) based on Beer-Lambert Law. NADH concentration vs. fluorescence units (FUs) standard curve ranging from 50 μM to 600 μM was measured at beginning of each measurement to determine the FUs to NADH concentration conversion factor. Raw fluorescence data from control reactions and kinase reactions for each condition were averaged and converted to picomole ADP produced using the NADH concentration conversion factor calculated assuming the amount of NDAH consumed is equivalent to the amount of ADP produced by ATP hydrolysis. Amount of ADP produced corrected for nonenzymatic ADP production (from control reactions) were plotted against time in Excel. The reaction rate of each experiment was calculated from the slope of the curve in the boxed regions.

### Kinase assay and western blotting

Kinase assays were performed by incubating 1 μM full-length BTK in a buffer containing 50 mM HEPES pH 7.4, 150 mM NaCl, 5% glycerol, 1mM DTT, 10 mM MgCl _2_ and 1 mM ATP. Activities of wild-type and mutant BTK proteins were carried out at room temperature. Samples from the reactions were collected and quenched at different time points by mixing with 4X SDS sample buffer for western blotting with the anti-pY551 antibody (BD Biosciences). The samples from 0 min were analyzed by SDS-PAGE as loading controls. Samples from 5, 10, 20 mins were analyzed by western blotting. The blots were quantified using the ChemiDoc detection system (BioRad).

### SAXS

Purified full-length BTK protein and the BTK SH3-SH2-kinase fragment (both with C-terminal hexa-His tags) at 10mg/ml were dialyzed against a buffer containing 201mM Tris, 1501mM NaCl, 1mM DTT and 2% glycerol. Four different concentrations of samples (1, 2, 5 and 10 mg/ml) were shipped to the MailinSAXS offered by the SIBYLS Beamline at the Advanced Light Source in Berkeley, California. The data was collected with X-ray wavelength at 1.271Å and the sample-to-detector distance at 21meters. Each sample was exposed to the beam for 10 sec with data collected as 0.2 sec slices, resulting total 50 frames of diffraction data per sample. Collected data were processed using the SIBYLS SAXS FrameSlice server (https://sibyls.als.lbl.gov/ran). ALMERGE was used to combine and extrapolate SAXS datasets collected at different concentrations to infinite dilution [68]. Merged data was analyzed with Raw [69]. Ab-initio reconstruction were performed with DAMMIF/N through Raw [70]. Reconstructed bead models were visualized with Chimera. Small Angle Scattering Biological Data Bank (SASBDB) accession codes: SASDRB9, SASDRC9, SASDRD9, SASDRE9.

## Acknowledgements

This work is supported by a grant from the National Institutes of Health (National Institute of Allergy and Infectious Diseases, AI43957) to A.H.A. We also thank the Roy J. Carver Charitable Trust, Muscatine, Iowa for ongoing research support and investment in the Iowa State University (ISU) Cryo-EM facility; data was collected on the Glacios 200kV TEM and Gatan K3 detector in this facility. Charles Stewart and the ISU Macromolecular X-ray Crystallography Facility arranged for shipping crystals and data collection. Greg Hura and the staff of the SIBYLS beamline at the Advanced Light Source provided training in SAXS data processing, interpretation, and data presentation for publication. ALS is a national user facility operated by Lawrence Berkeley National Laboratory on behalf of the Department of Energy, Office of Basic Energy Sciences, through the Integrated Diffraction Analysis Technologies (IDAT) program, supported by DOE Office of Biological and Environmental Research. Additional support comes from the National Institute of Health project ALS-ENABLE (P30 GM124169) and a High-End Instrumentation Grant S10OD018483. This research also used resources of GM/CA and NE-CAT beamlines at the Advanced Photon Source. GM/CA has been funded by the National Cancer Institute (ACB-12002), the National Institute of General Medical Sciences (AGM-12006, P30GM138396) and the NIH shared instrumentation grant (S10 OD012289). NECAT has been funded by the National Institutes of Health (P30 GM124165) and a NIH-ORIP HEI grant (S10OD021527).

## Data availability

Coordinates and structure factors for Btk crystal structures are deposited in the PDB; cryoEM maps are deposited in the EMDB; SAXS data are deposited in the SASBDB; Hydrogen/deuterium exchange data are deposited in the PRIDE database.

---

**Fig. 2.**
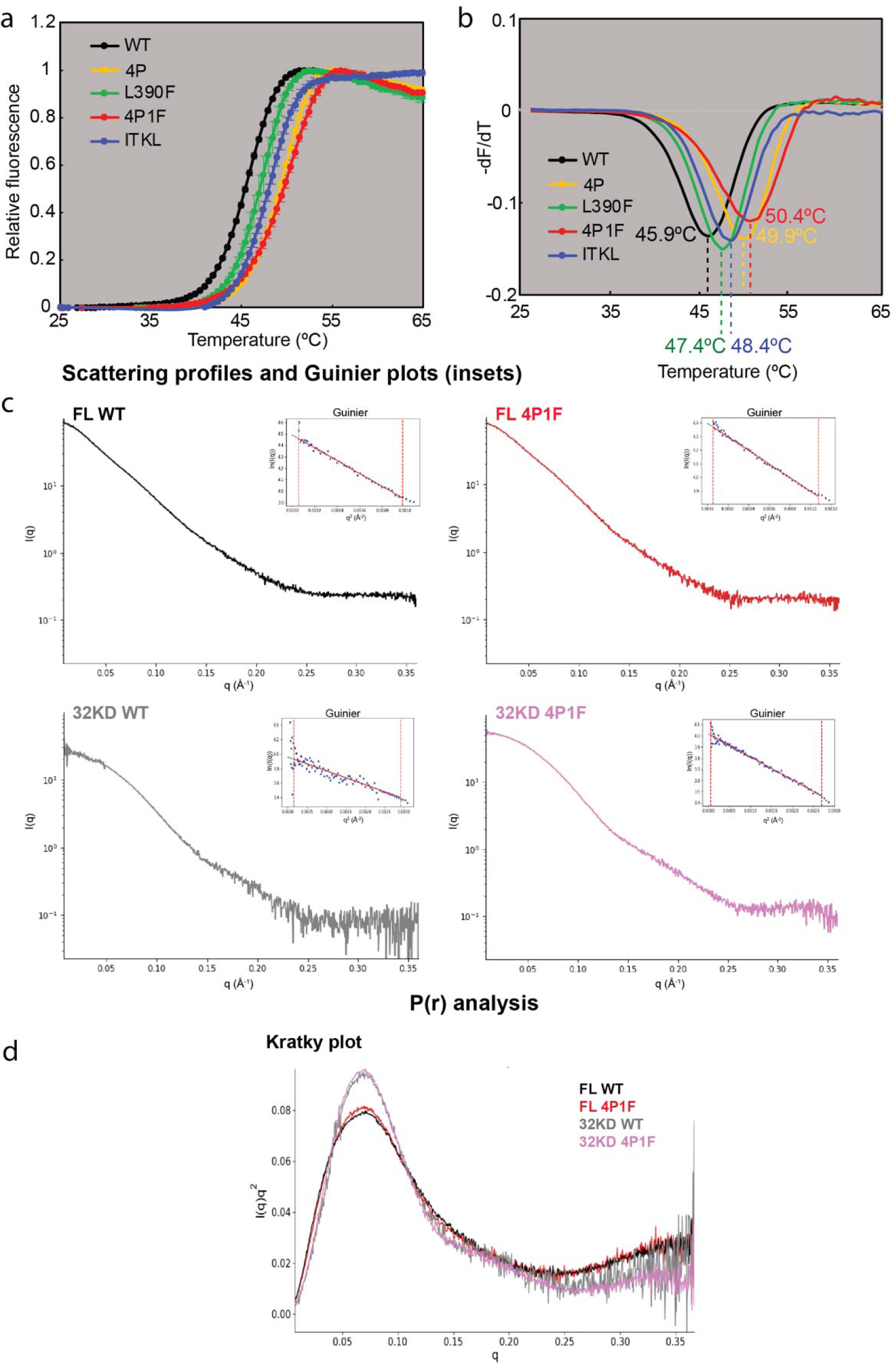
Figure supplement 1. (a) T_m_ curves and (b) first derivatives for panel of BTK variants. (c) Guinier and (d) Kratky plots for BTK variants.

**Fig. 4.**
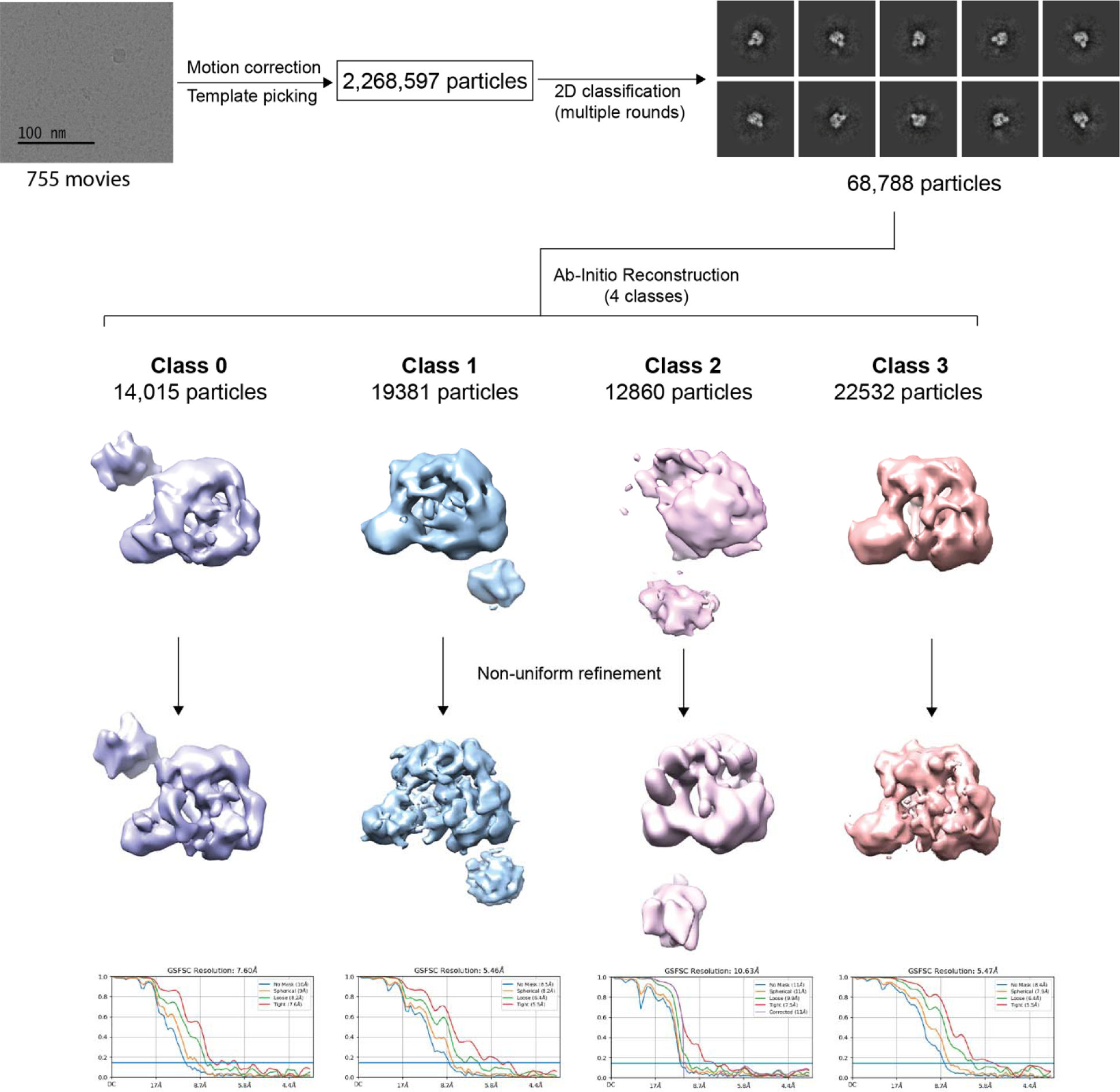
Figure supplement 1. Workflow showing cryoEM analysis of full-length BTK.

**Fig. 4.**
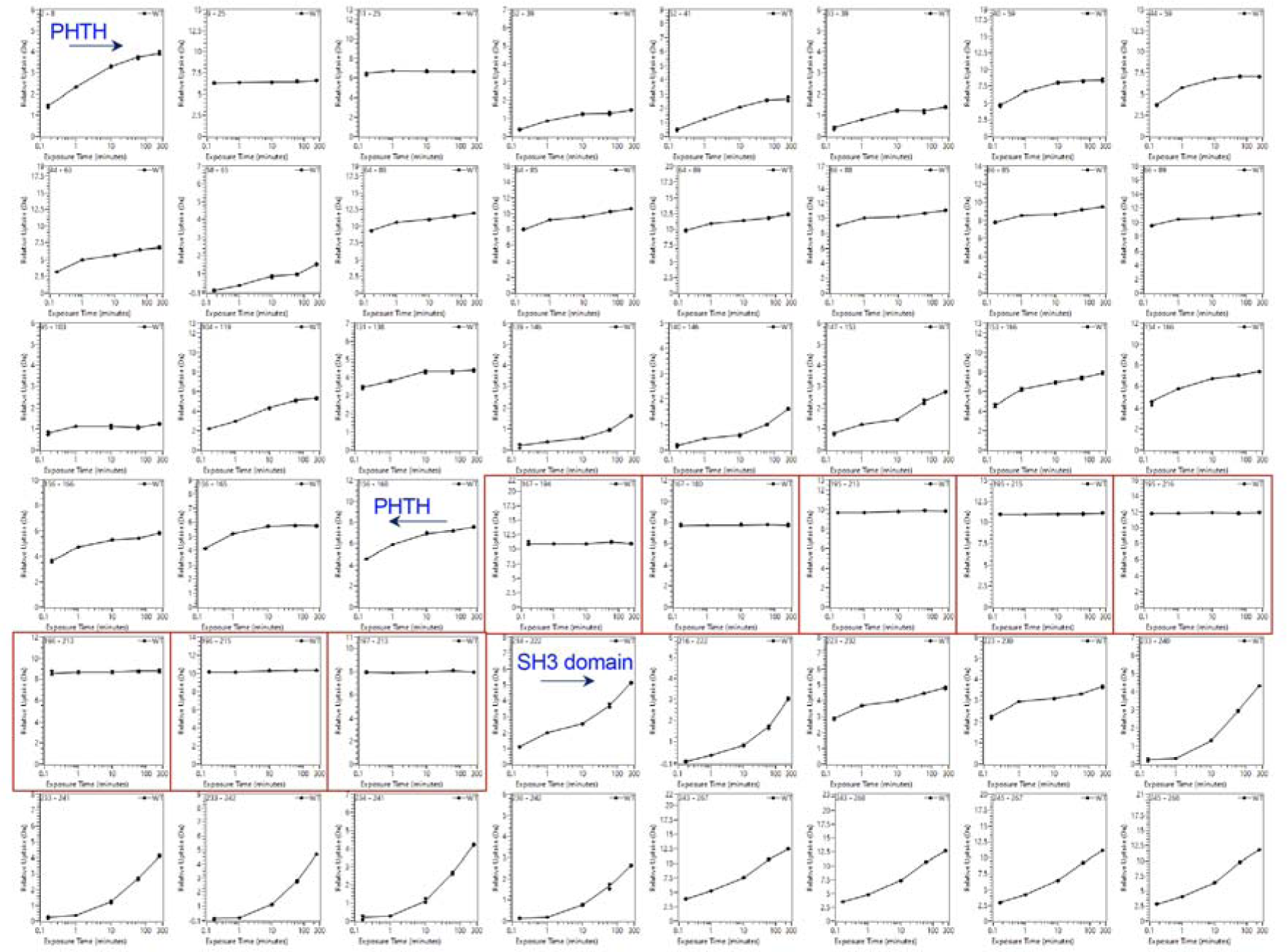
Figure supplement 2. Deuterium uptake curves for the PHTH through SH3 domain of BTK. Domain boundaries for the PHTH and the start of the SH3 domain are indicated. Uptake curves for peptides derived from the linker between PHTH and SH3 (residues 171-214) are boxed in red. The complete HDX dataset is provided in the supplemental data file.

## References

1. Yu L, Smith CI. Tec family kinases. FEBS J. 2011;278(12):1969. Epub 2011/04/27. doi: 10.1111/j.1742-4658.2011.08135.x. PubMed PMID: 21518256.

2. Mano H. Tec family of protein-tyrosine kinases: an overview of their structure and function. Cytokine Growth Factor Rev. 1999;10(3-4):267–80. Epub 2000/01/27. doi: 10.1016/s1359-6101(99)00019-2. PubMed PMID: 10647781.

3. Smith CI, Islam TC, Mattsson PT, Mohamed AJ, Nore BF, Vihinen M. The Tec family of cytoplasmic tyrosine kinases: mammalian Btk, Bmx, Itk, Tec, Txk and homologs in other species. Bioessays. 2001;23(5):436–46. Epub 2001/05/08. doi: 10.1002/bies.1062. PubMed PMID: 11340625.

4. Xu W, Harrison SC, Eck MJ. Three-dimensional structure of the tyrosine kinase c-Src. Nature. 1997;385(6617):595–602. PubMed PMID: 9024657.

5. Wang Q, Vogan EM, Nocka LM, Rosen CE, Zorn JA, Harrison SC, et al. Autoinhibition of Bruton’s tyrosine kinase (Btk) and activation by soluble inositol hexakisphosphate. Elife. 2015;4. Epub 2015/02/24. doi: 10.7554/eLife.06074. PubMed PMID: 25699547; PubMed Central PMCID: PMCPMC4384635.

6. Joseph RE, Wales TE, Fulton DB, Engen JR, Andreotti AH. Achieving a Graded Immune Response: BTK Adopts a Range of Active/Inactive Conformations Dictated by Multiple Interdomain Contacts. Structure. 2017;25(10):1481–94 e4. Epub 2017/09/05. doi: 10.1016/j.str.2017.07.014. PubMed PMID: 28867612; PubMed Central PMCID: PMCPMC5629114.

7. Koegl M, Zlatkine P, Ley SC, Courtneidge SA, Magee AI. Palmitoylation of multiple Src-family kinases at a homologous N-terminal motif. Biochem J. 1994;303 (Pt 3)(Pt 3):749–53. Epub 1994/11/01. doi: 10.1042/bj3030749. PubMed PMID: 7980442; PubMed Central PMCID: PMCPMC1137610.

8. Baraldi E, Carugo KD, Hyvonen M, Surdo PL, Riley AM, Potter BV, et al. Structure of the PH domain from Bruton’s tyrosine kinase in complex with inositol 1,3,4,5-tetrakisphosphate. Structure. 1999;7(4):449–60. PubMed PMID: 10196129.

9. Hyvonen M, Saraste M. Structure of the PH domain and Btk motif from Bruton’s tyrosine kinase: molecular explanations for X-linked agammaglobulinaemia. Embo J. 1997;16(12):3396–404. PubMed PMID: 9218782.

10. Devkota S, Joseph RE, Boyken SE, Fulton DB, Andreotti AH. An Autoinhibitory Role for the Pleckstrin Homology Domain of Interleukin-2-Inducible Tyrosine Kinase and Its Interplay with Canonical Phospholipid Recognition. Biochemistry. 2017;56(23):2938–49. Epub 2017/05/19. doi: 10.1021/acs.biochem.6b01182. PubMed PMID: 28516764; PubMed Central PMCID: PMCPMC5956896.

11. Amatya N, Wales TE, Kwon A, Yeung W, Joseph RE, Fulton DB, et al. Lipid-targeting pleckstrin homology domain turns its autoinhibitory face toward the TEC kinases. Proc Natl Acad Sci U S A. 2019;116(43):21539–44. Epub 2019/10/09. doi: 10.1073/pnas.1907566116. PubMed PMID: 31591208; PubMed Central PMCID: PMCPMC6815127.

12. Lerner EC, Trible RP, Schiavone AP, Hochrein JM, Engen JR, Smithgall TE. Activation of the Src family kinase Hck without SH3-linker release. J Biol Chem. 2005;280(49):40832–7. Epub 2005/10/08. doi: 10.1074/jbc.M508782200. PubMed PMID: 16210316.

13. von Raussendorf F, de Ruiter A, Leonard TA. A switch in nucleotide affinity governs activation of the Src and Tec family kinases. Sci Rep. 2017;7(1):17405. Epub 2017/12/14. doi: 10.1038/s41598-017-17703-5. PubMed PMID: 29234112; PubMed Central PMCID: PMCPMC5727165.

14. Joseph RE, Kleino I, Wales TE, Xie Q, Fulton DB, Engen JR, et al. Activation loop dynamics determine the different catalytic efficiencies of B cell- and T cell-specific tec kinases. Sci Signal. 2013;6(290):ra76. Epub 2013/08/29. doi: 10.1126/scisignal.2004298. PubMed PMID: 23982207; PubMed Central PMCID: PMCPMC4089953.

15. Duarte DP, Lamontanara AJ, La Sala G, Jeong S, Sohn YK, Panjkovich A, et al. Btk SH2-kinase interface is critical for allosteric kinase activation and its targeting inhibits B-cell neoplasms. Nat Commun. 2020;11(1):2319. Epub 2020/05/10. doi: 10.1038/s41467-020-16128-5. PubMed PMID: 32385234; PubMed Central PMCID: PMCPMC7210950.

16. Marquez JA, Smith CI, Petoukhov MV, Lo Surdo P, Mattsson PT, Knekt M, et al. Conformation of full-length Bruton tyrosine kinase (Btk) from synchrotron X-ray solution scattering. Embo J. 2003;22(18):4616–24. PubMed PMID: 12970174.

17. Goldschmidt L, Cooper DR, Derewenda ZS, Eisenberg D. Toward rational protein crystallization: A Web server for the design of crystallizable protein variants. Protein Sci. 2007;16(8):1569–76. Epub 2007/07/28. doi: 10.1110/ps.072914007. PubMed PMID: 17656576; PubMed Central PMCID: PMCPMC2203352.

18. Gonfloni S, Williams JC, Hattula K, Weijland A, Wierenga RK, Superti-Furga G. The role of the linker between the SH2 domain and catalytic domain in the regulation and function of Src. EMBO J. 1997;16(24):7261–71. Epub 1998/02/21. doi: 10.1093/emboj/16.24.7261. PubMed PMID: 9405355; PubMed Central PMCID: PMCPMC1170326.

19. Dinh M, Grunberger D, Ho H, Tsing SY, Shaw D, Lee S, et al. Activation mechanism and steady state kinetics of Bruton’s tyrosine kinase. J Biol Chem. 2007;282(12):8768–76. Epub 2007/02/01. doi: 10.1074/jbc.M609920200. PubMed PMID: 17264076.

20. Chung JK, Nocka LM, Decker A, Wang Q, Kadlecek TA, Weiss A, et al. Switch-like activation of Bruton’s tyrosine kinase by membrane-mediated dimerization. Proc Natl Acad Sci U S A. 2019;116(22):10798–803. Epub 2019/05/12. doi: 10.1073/pnas.1819309116. PubMed PMID: 31076553; PubMed Central PMCID: PMCPMC6561188.

21. Joseph RE, Amatya N, Fulton DB, Engen JR, Wales TE, Andreotti A. Differential impact of BTK active site inhibitors on the conformational state of full-length BTK. Elife. 2020;9. Epub 2020/11/24. doi: 10.7554/eLife.60470. PubMed PMID: 33226337; PubMed Central PMCID: PMCPMC7834017.

22. Marcotte DJ, Liu YT, Arduini RM, Hession CA, Miatkowski K, Wildes CP, et al. Structures of human Bruton’s tyrosine kinase in active and inactive conformations suggest a mechanism of activation for TEC family kinases. Protein Sci. 2010;19(3):429–39. Epub 2010/01/07. doi: 10.1002/pro.321. PubMed PMID: 20052711; PubMed Central PMCID: PMCPMC2866269.

23. Beenstock J, Mooshayef N, Engelberg D. How Do Protein Kinases Take a Selfie (Autophosphorylate)? Trends Biochem Sci. 2016;41(11):938–53. Epub 2016/10/30. doi: 10.1016/j.tibs.2016.08.006. PubMed PMID: 27594179.

24. Dodson CA, Kosmopoulou M, Richards MW, Atrash B, Bavetsias V, Blagg J, et al. Crystal structure of an Aurora-A mutant that mimics Aurora-B bound to MLN8054: insights into selectivity and drug design. Biochem J. 2010;427(1):19–28. Epub 2010/01/14. doi: 10.1042/BJ20091530. PubMed PMID: 20067443.

25. Colombano G, Caldwell JJ, Matthews TP, Bhatia C, Joshi A, McHardy T, et al. Binding to an Unusual Inactive Kinase Conformation by Highly Selective Inhibitors of Inositol-Requiring Enzyme 1alpha Kinase-Endoribonuclease. J Med Chem. 2019;62(5):2447–65. Epub 2019/02/20. doi: 10.1021/acs.jmedchem.8b01721. PubMed PMID: 30779566; PubMed Central PMCID: PMCPMC6437697.

26. Modi V, Dunbrack RL, Jr. Defining a new nomenclature for the structures of active and inactive kinases. Proc Natl Acad Sci U S A. 2019;116(14):6818–27. Epub 2019/03/15. doi: 10.1073/pnas.1814279116. PubMed PMID: 30867294; PubMed Central PMCID: PMCPMC6452665.

27. Joseph RE, Xie Q, Andreotti AH. Identification of an allosteric signaling network within Tec family kinases. J Mol Biol. 2010;403(2):231–42. Epub 2010/09/10. doi: 10.1016/j.jmb.2010.08.035. PubMed PMID: 20826165; PubMed Central PMCID: PMCPMC2949508.

28. Muckelbauer J, Sack JS, Ahmed N, Burke J, Chang CY, Gao M, et al. X-ray crystal structure of bone marrow kinase in the x chromosome: a Tec family kinase. Chem Biol Drug Des. 2011;78(5):739–48. Epub 2011/09/03. doi: 10.1111/j.1747-0285.2011.01230.x. PubMed PMID: 21883956.

29. Garland-Kuntz EE, Vago FS, Sieng M, Van Camp M, Chakravarthy S, Blaine A, et al. Direct observation of conformational dynamics of the PH domain in phospholipases Cl’. and beta may contribute to subfamily-specific roles in regulation. J Biol Chem. 2018;293(45):17477–90. Epub 2018/09/23. doi: 10.1074/jbc.RA118.003656. PubMed PMID: 30242131; PubMed Central PMCID: PMCPMC6231117.

30. Ashwell MA, Lapierre JM, Brassard C, Bresciano K, Bull C, Cornell-Kennon S, et al. Discovery and optimization of a series of 3-(3-phenyl-3H-imidazo[4,5-b]pyridin-2-yl)pyridin-2-amines: orally bioavailable, selective, and potent ATP-independent Akt inhibitors. J Med Chem. 2012;55(11):5291–310. Epub 2012/04/27. doi: 10.1021/jm300276x. PubMed PMID: 22533986.

31. Quambusch L, Landel I, Depta L, Weisner J, Uhlenbrock N, Muller MP, et al. Covalent-Allosteric Inhibitors to Achieve Akt Isoform-Selectivity. Angew Chem Int Ed Engl. 2019;58(52):18823–9. Epub 2019/10/05. doi: 10.1002/anie.201909857. PubMed PMID: 31584233; PubMed Central PMCID: PMCPMC6972997.

32. Truebestein L, Hornegger H, Anrather D, Hartl M, Fleming KD, Stariha JTB, et al. Structure of autoinhibited Akt1 reveals mechanism of PIP(3)-mediated activation. Proc Natl Acad Sci U S A. 2021;118(33). Epub 2021/08/14. doi: 10.1073/pnas.2101496118. PubMed PMID: 34385319; PubMed Central PMCID: PMCPMC8379990.

33. Wu WI, Voegtli WC, Sturgis HL, Dizon FP, Vigers GP, Brandhuber BJ. Crystal structure of human AKT1 with an allosteric inhibitor reveals a new mode of kinase inhibition. PLoS One. 2010;5(9):e12913. Epub 2010/10/05. doi: 10.1371/journal.pone.0012913. PubMed PMID:

34. 20886116; PubMed Central PMCID: PMCPMC2944833

34. Bae H, Viennet T, Park E, Chu N, Salguero A, Eck MJ, et al. PH domain-mediated autoinhibition and oncogenic activation of Akt. Elife. 2022;11. Epub 2022/08/16. doi: 10.7554/eLife.80148. PubMed PMID: 35968932; PubMed Central PMCID: PMCPMC9417420.

35. Shaw AL, Parson MAH, Truebestein L, Jenkins ML, Leonard TA, Burke JE. ATP-competitive and allosteric inhibitors induce differential conformational changes at the autoinhibitory interface of Akt1. Structure. 2023;31(3):343–54 e3. Epub 2023/02/10. doi: 10.1016/j.str.2023.01.007. PubMed PMID: 36758543.

36. Arbesu M, Maffei M, Cordeiro TN, Teixeira JM, Perez Y, Bernado P, et al. The Unique Domain Forms a Fuzzy Intramolecular Complex in Src Family Kinases. Structure. 2017;25(4):630–40 e4. Epub 2017/03/21. doi: 10.1016/j.str.2017.02.011. PubMed PMID: 28319009.

37. Perez Y, Maffei M, Igea A, Amata I, Gairi M, Nebreda AR, et al. Lipid binding by the Unique and SH3 domains of c-Src suggests a new regulatory mechanism. Sci Rep. 2013;3:1295. Epub 2013/02/19. doi: 10.1038/srep01295. PubMed PMID: 23416516; PubMed Central PMCID: PMCPMC3575015.

38. Pluk H, Dorey K, Superti-Furga G. Autoinhibition of c-Abl. Cell. 2002;108(2):247-59. Epub 2002/02/08. doi: 10.1016/s0092-8674(02)00623-2. PubMed PMID: 11832214.

39. Nagar B, Hantschel O, Young MA, Scheffzek K, Veach D, Bornmann W, et al. Structural basis for the autoinhibition of c-Abl tyrosine kinase. Cell. 2003;112(6):859–71. Epub 2003/03/26. doi: 10.1016/s0092-8674(03)00194-6. PubMed PMID: 12654251.

40. Hantschel O, Nagar B, Guettler S, Kretzschmar J, Dorey K, Kuriyan J, et al. A myristoyl/phosphotyrosine switch regulates c-Abl. Cell. 2003;112(6):845–57. Epub 2003/03/26. doi: 10.1016/s0092-8674(03)00191-0. PubMed PMID: 12654250.

41. Panjarian S, Iacob RE, Chen S, Engen JR, Smithgall TE. Structure and dynamic regulation of Abl kinases. J Biol Chem. 2013;288(8):5443–50. Epub 2013/01/15. doi: 10.1074/jbc.R112.438382. PubMed PMID: 23316053; PubMed Central PMCID: PMCPMC3581414.

42. Laederach A, Cradic KW, Brazin KN, Zamoon J, Fulton DB, Huang XY, et al. Competing modes of self-association in the regulatory domains of Bruton’s tyrosine kinase: intramolecular contact versus asymmetric homodimerization. Protein Sci. 2002;11(1):36–45. PubMed PMID: 11742120.

43. Liu W, Quinto I, Chen X, Palmieri C, Rabin RL, Schwartz OM, et al. Direct inhibition of Bruton’s tyrosine kinase by IBtk, a Btk-binding protein. Nat Immunol. 2001;2(10):939–46. Epub 2001/09/29. doi: 10.1038/ni1001-939. PubMed PMID: 11577348.

44. Nocka LM, Groves, J.T., Kuriyan, J. Stimulation of the catalytic activity of the tyrosine kinase Btk by the adaptor protein Grb2. 2023. doi: 10.1101/2022.08.25.505243.

45. Tsukada S, Simon MI, Witte ON, Katz A. Binding of beta gamma subunits of heterotrimeric G proteins to the PH domain of Bruton tyrosine kinase. Proc Natl Acad Sci U S A. 1994;91(23):11256–60. Epub 1994/11/08. doi: 10.1073/pnas.91.23.11256. PubMed PMID: 7972043; PubMed Central PMCID: PMCPMC45206.

46. Shelby SA, Castello-Serrano I, Wisser KC, Levental I, Veatch SL. Membrane phase separation drives responsive assembly of receptor signaling domains. Nat Chem Biol. 2023. Epub 2023/03/31. doi: 10.1038/s41589-023-01268-8. PubMed PMID: 36997644.

47. Nocka L, Groves, J.T., Kuriyan, J. Stimulation of the catalytic activity of the tyrosine kinase Btk by the adaptor protein Grb2. bioRxiv 20220825505243. 2022. doi: 10.1101/2022.08.25.505243.

48. Grassilli E, Pisano F, Cialdella A, Bonomo S, Missaglia C, Cerrito MG, et al. A novel oncogenic BTK isoform is overexpressed in colon cancers and required for RAS-mediated transformation. Oncogene. 2016;35(33):4368–78. Epub 2016/01/26. doi: 10.1038/onc.2015.504. PubMed PMID: 26804170; PubMed Central PMCID: PMCPMC4994017.

49. Eifert C, Wang X, Kokabee L, Kourtidis A, Jain R, Gerdes MJ, et al. A novel isoform of the B cell tyrosine kinase BTK protects breast cancer cells from apoptosis. Genes Chromosomes Cancer. 2013;52(10):961–75. Epub 2013/08/06. doi: 10.1002/gcc.22091. PubMed PMID: 23913792; PubMed Central PMCID: PMCPMC5006942.

50. Wang X, Kokabee L, Kokabee M, Conklin DS. Bruton’s Tyrosine Kinase and Its Isoforms in Cancer. Front Cell Dev Biol. 2021;9:668996. Epub 2021/07/27. doi: 10.3389/fcell.2021.668996. PubMed PMID: 34307353; PubMed Central PMCID: PMCPMC8297165.

51. Grassilli E, Cerrito MG, Lavitrano M. BTK, the new kid on the (oncology) block? Front Oncol. 2022;12:944538. Epub 2022/08/23. doi: 10.3389/fonc.2022.944538. PubMed PMID: 35992808; PubMed Central PMCID: PMCPMC9386470.

52. Shah NH, Amacher JF, Nocka LM, Kuriyan J. The Src module: an ancient scaffold in the evolution of cytoplasmic tyrosine kinases. Crit Rev Biochem Mol Biol. 2018;53(5):535–63. Epub 2018/09/06. doi: 10.1080/10409238.2018.1495173. PubMed PMID: 30183386; PubMed Central PMCID: PMCPMC6328253.

53. Vonrhein C, Flensburg C, Keller P, Sharff A, Smart O, Paciorek W, et al. Data processing and analysis with the autoPROC toolbox. Acta Crystallogr D Biol Crystallogr. 2011;67(Pt 4):293–302. Epub 2011/04/05. doi: 10.1107/S0907444911007773. PubMed PMID: 21460447; PubMed Central PMCID: PMCPMC3069744.

54. Tickle IJ, Flensburg, C., Keller, P., Paciorek, W., Sharff, A.,, Vonrhein C, Bricogne, G. Staraniso. Cambridge, United Kingdom: Global Phasing Ltd. 2018.

55. Winn MD, Ballard CC, Cowtan KD, Dodson EJ, Emsley P, Evans PR, et al. Overview of the CCP4 suite and current developments. Acta Crystallogr D Biol Crystallogr. 2011;67(Pt 4):235–42. Epub 2011/04/05. doi: 10.1107/S0907444910045749. PubMed PMID: 21460441; PubMed Central PMCID: PMCPMC3069738.

56. McCoy AJ, Grosse-Kunstleve RW, Adams PD, Winn MD, Storoni LC, Read RJ. Phaser crystallographic software. J Appl Crystallogr. 2007;40(Pt 4):658-74. Epub 2007/08/01. doi: 10.1107/S0021889807021206. PubMed PMID: 19461840; PubMed Central PMCID: PMCPMC2483472.

57. Liebschner D, Afonine PV, Baker ML, Bunkoczi G, Chen VB, Croll TI, et al. Macromolecular structure determination using X-rays, neutrons and electrons: recent developments in Phenix. Acta Crystallogr D Struct Biol. 2019;75(Pt 10):861–77. Epub 2019/10/08. doi: 10.1107/S2059798319011471. PubMed PMID: 31588918; PubMed Central PMCID: PMCPMC6778852.

58. Emsley P, Cowtan K. Coot: model-building tools for molecular graphics. Acta Crystallogr D Biol Crystallogr. 2004;60(Pt 12 Pt 1):2126–32. Epub 2004/12/02. doi: 10.1107/S0907444904019158. PubMed PMID: 15572765.

59. DeLano WL. The PyMOL Molecular Graphics System DeLano Scientific, San Carlos, CA, USA. 2002.

60. Crawford JJ, Johnson AR, Misner DL, Belmont LD, Castanedo G, Choy R, et al. Discovery of GDC-0853: A Potent, Selective, and Noncovalent Bruton’s Tyrosine Kinase Inhibitor in Early Clinical Development. J Med Chem. 2018;61(6):2227–45. Epub 2018/02/20. doi: 10.1021/acs.jmedchem.7b01712. PubMed PMID: 29457982.

61. Punjani A, Rubinstein JL, Fleet DJ, Brubaker MA. cryoSPARC: algorithms for rapid unsupervised cryo-EM structure determination. Nat Methods. 2017;14(3):290–6. Epub 2017/02/07. doi: 10.1038/nmeth.4169. PubMed PMID: 28165473.

62. Punjani A, Zhang H, Fleet DJ. Non-uniform refinement: adaptive regularization improves single-particle cryo-EM reconstruction. Nat Methods. 2020;17(12):1214–21. Epub 2020/12/02. doi: 10.1038/s41592-020-00990-8. PubMed PMID: 33257830.

63. Pettersen EF, Goddard TD, Huang CC, Couch GS, Greenblatt DM, Meng EC, et al. UCSF Chimera--a visualization system for exploratory research and analysis. J Comput Chem. 2004;25(13):1605–12. Epub 2004/07/21. doi: 10.1002/jcc.20084. PubMed PMID: 15264254.

64. Wales TE, Engen JR. Hydrogen exchange mass spectrometry for the analysis of protein dynamics. Mass Spectrom Rev. 2006;25(1):158–70. Epub 2005/10/07. doi: 10.1002/mas.20064. PubMed PMID: 16208684.

65. Masson GR, Burke JE, Ahn NG, Anand GS, Borchers C, Brier S, et al. Recommendations for performing, interpreting and reporting hydrogen deuterium exchange mass spectrometry (HDX-MS) experiments. Nat Methods. 2019;16(7):595–602. Epub 2019/06/30. doi: 10.1038/s41592-019-0459-y. PubMed PMID: 31249422; PubMed Central PMCID: PMCPMC6614034.

66. Perez-Riverol Y, Bai J, Bandla C, Garcia-Seisdedos D, Hewapathirana S, Kamatchinathan S, et al. The PRIDE database resources in 2022: a hub for mass spectrometry-based proteomics evidences. Nucleic Acids Res. 2022;50(D1):D543–D52. Epub 2021/11/02. doi: 10.1093/nar/gkab1038. PubMed PMID: 34723319; PubMed Central PMCID: PMCPMC8728295.

67. Barker SC, Kassel DB, Weigl D, Huang X, Luther MA, Knight WB. Characterization of pp60c-src tyrosine kinase activities using a continuous assay: autoactivation of the enzyme is an intermolecular autophosphorylation process. Biochemistry. 1995;34(45):14843–51. Epub 1995/11/14. doi: 10.1021/bi00045a027. PubMed PMID: 7578094.

68. Franke D, Kikhney, A.G., Svergun, D.I. Automated acquisition and analysis of small angle X-ray scattering data. Nuclear Instruments and Methods in Physics Research - section A. 2012;689:52–9. doi: 10.1016/j.nima.2012.06.008.

69. Hopkins JB, Gillilan, R.E., Skou, S. BioXTAS RAW: improvements to a free open-source program for small-angle X-ray scattering data reduction and analysis. J Appl Cryst. 2017;50:1545–53. doi: 10.1107/S1600576717011438.

70. Franke D, Svergun, D.I. DAMMIF, a program for rapid ab-initio shape determination in small-angle scattering. J Appl Cryst. 2009;42:342–6. doi: 10.1107/S0021889809000338.

71. Wriggers W. Conventions and workflows for using Situs. Acta Crystallogr D Biol Crystallogr. 2012;68(Pt 4):344–51. Epub 2012/04/17. doi: 10.1107/S0907444911049791. PubMed PMID: 22505255; PubMed Central PMCID: PMCPMC3322594.

72. Chopra N, Wales TE, Joseph RE, Boyken SE, Engen JR, Jernigan RL, et al. Dynamic Allostery Mediated by a Conserved Tryptophan in the Tec Family Kinases. PLoS Comput Biol. 2016;12(3):e1004826. Epub 2016/03/25. doi: 10.1371/journal.pcbi.1004826. PubMed PMID: 27010561; PubMed Central PMCID: PMCPMC4807093.

